# Developing Anti-EGFR/Anti-HER2 Bifunctional Antibody for Solid Tumors by Protein Engineering

**DOI:** 10.64898/2025.12.25.696462

**Authors:** Senem Sen, Aslı Semerci, Melis Karaca, Recep Erdem Ahan, Cemile Elif Ozcelik, Elif Duman, Behide Saltepe, Ebru Sahin Kehribar, Ebru Aras, Eray Ulas Bozkurt, Hilal Yazıcı, Müge Serhatlı, Urartu Ozgur Safak Seker

## Abstract

Overexpression of epidermal growth factor receptor (EGFR) and human epidermal growth factor receptor 2 (HER2) is common in solid tumors like breast, colorectal, and head and neck cancers, driving oncogenic signaling, therapy resistance, and poor prognosis. Monotherapies often fall short in fully suppressing tumor progression, particularly in cases with EGFR/HER2 co-amplification. Dual-targeting strategies offer enhanced efficacy by mitigating resistance and amplifying antitumor responses. Our study focused on developing and characterizing a bispecific anti-EGFR/HER2 antibody designed to simultaneously block ligand binding and receptor activation while harnessing Fc-mediated immune effector functions. Utilizing knob-into-hole and CrossMab technologies, we engineered a bispecific antibody capable of binding the extracellular domains of both EGFR and HER2. *In vitro* analyses confirmed its dual-binding capacity. Comparative studies revealed that this bispecific construct outperformed its monospecific counterparts in suppressing proliferation and, notably, in increasing the expression of apoptotic markers in EGFR/HER2-expressing tumor cells. Together, these findings highlight the synergistic therapeutic potential of bispecific antibodies as a promising modality for overcoming the limitations of current monotherapies.

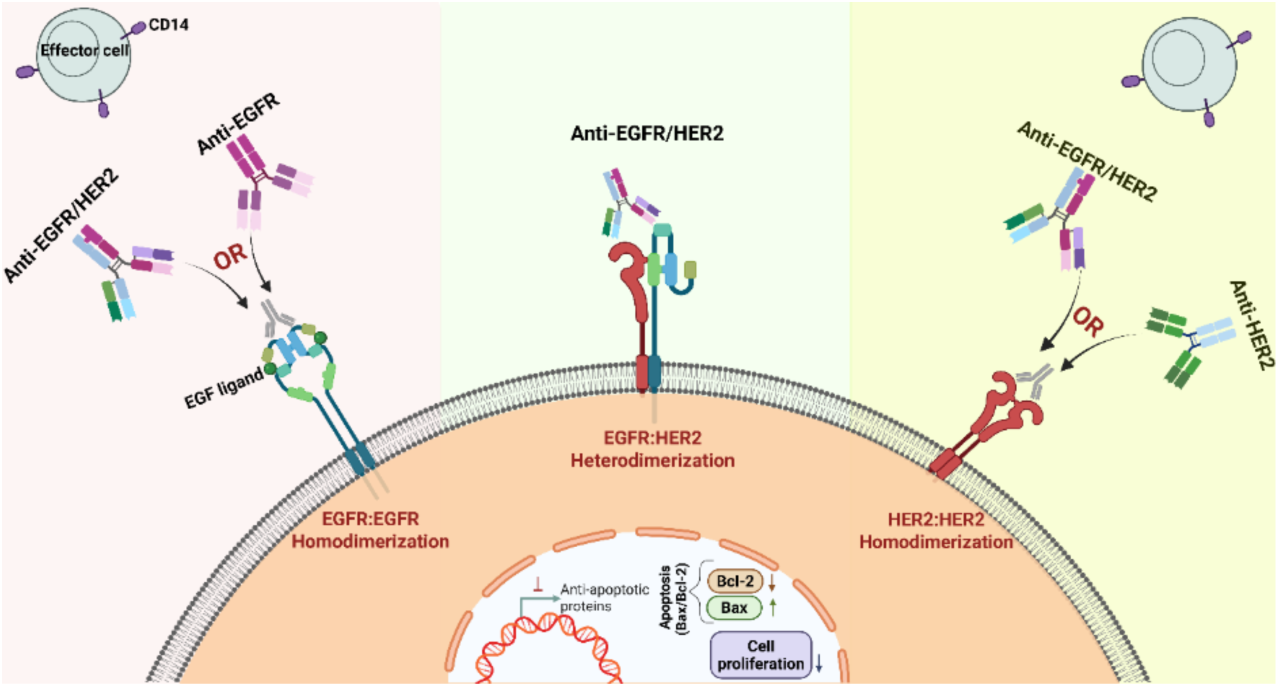

## INTRODUCTION

A major challenge in oncology is the selective eradication of malignant cells while minimizing collateral damage to normal tissues. Although traditional therapies have saved countless lives, they often cause significant toxicity, highlighting the need for more precise treatment strategies. Targeted therapies have emerged to fill this gap, demonstrating improved therapeutic indices compared to conventional approaches. ^1^ Despite advances in early diagnosis and intervention, issues such as drug resistance, metastasis, and recurrence continue to compromise long-term patient outcomes.^2^

Cancer cells follow complex pathways to survive and proliferate, necessitating more specific treatment approaches. Monoclonal antibodies (mAbs) have revolutionized cancer therapy, and since their first approval in 1986, they have become a central component in the management of both immune disorders and malignancies.^3^

As our understanding of cancer biology deepens, immunotherapy strategies are evolving to maintain their effectiveness. This evolution has led to the development of novel molecular formats extending beyond traditional mAbs. These include fragment-based bispecific antibodies (BsAbs), Fc-based formats mimicking native IgG, immune checkpoint inhibitors, antibody-drug conjugates, CAR-modified immune cells, and smaller constructs such as dual-affinity retargeting molecules (DARTs®), bispecific T-cell engagers (BiTEs®), bispecific killer cell engagers (BiKEs®), trispecific killer cell engagers (TriKEs®), nanobodies, and tandem diabodies (TandAbs®). All are designed to improve targeting, stability, and therapeutic delivery.^2,4–6^

Through molecular optimization, these agents leverage antibody specificity to orchestrate complex immune responses against tumors.^6,7^ Notable milestones include the approval of trastuzumab as the first mAb therapy for solid tumors and blinatumomab as the first clinically successful bispecific antibody.^8^ Further advancements in antibody engineering, such as CrossMab and knobs-into-holes (KIH) technologies, have resolved key challenges in bispecific design by improving heterodimer pairing and enhancing functional stability.^9–12^ These innovations collectively mark a paradigm shift toward more targeted, multifunctional, and adaptable cancer immunotherapies.

The epidermal growth factor receptor (EGFR) family is critically involved in the pathogenesis of various cancers. Targeted therapies, including monoclonal antibodies and tyrosine kinase inhibitors (TKIs), have been developed to address malignancies such as HER2-positive breast cancer and EGFR-expressing colorectal and head-and-neck cancers.^13–15^ Early discoveries in the 1980s highlighted the structural homology between EGFR and HER2, predicting ligand-induced ERBB2 activation.^16^ Subsequent studies confirmed EGFR–HER2 heterodimerization, enhancing oncogenic signaling.^17^ When it comes to treatment strategies, individuals with EGFR and HER2 co-amplification display resistance to conventional therapies such as chemotherapy, radiotherapy, and hormonal treatments. This suggests the need for tailored therapeutic approaches for this specific subgroup of patients.^18^

Success with monotherapies using engineered antibodies in therapeutic systems is strong but limited. Consequently, the integration of combination strategies has emerged as a crucial advancement in improving therapeutic outcomes. Dual-targeting approaches that simultaneously engage two antigens overcome limitations of traditional monotherapies by offering enhanced antitumor activity and synergistic therapeutic responses.^19,20^ In particular, the limited success of ErbB signaling inhibitors as single agents highlights the need for co-targeting both EGFR and HER2 pathways. Clinical studies combining Cetuximab and Trastuzumab with Taxotere-based chemotherapy have shown improved elimination of tumor-initiating cells and better clinical outcomes.^20^ Supporting this, preclinical models revealed that combining Osimertinib, Cetuximab, and Trastuzumab effectively overcame resistance mechanisms in lung cancer.^21^ Dual use of EGFR-targeting antibodies like cetuximab and trastuzumab has also been shown to inhibit angiogenesis, cell proliferation, and downstream signaling more effectively than either agent alone.^22^ Dual-specificity inhibitors, such as lapatinib, neratinib, and afatinib, now represent therapeutic options targeting both the EGFR and HER2 pathways. For example, lapatinib —a dual EGFR/ERBB2 tyrosine kinase inhibitor—has been clinically approved. However, its activity may be diminished by human serum factors that reverse cell cycle arrest and restore oncogenic signaling.^23^ Unlike these small molecules that block intracellular kinase activity, bispecific antibodies bind EGFR and HER2 extracellularly, blocking ligands and internalizing receptors, initiating immune-mediated effects. Recent single-molecule imaging studies revealed the dynamic nature of EGFR and HER2 interactions, including pre-formed receptor clusters and transient dimerization, which supports the rationale for dual-targeting.^24^ Antibodies that promote HER2/HER2 and HER2/EGFR internalization are particularly promising for overcoming resistance.^25^ Dual-targeted therapies have been shown to have superior tumor regression and survival benefits compared to monotherapies, suggesting that co-targeting these receptors may effectively address the limitations associated with single-agent treatments.^26^

Given the complexity of signaling and resistance in cancer, combining bifunctional agents that block multiple pathways is essential. Combining different anti-receptor antibodies can inhibit a cancer model driven by EGFR and also increase antitumor efficacy.

In this study, we hypothesized that an anti-EGFR/HER2 bispecific antibody, which blocks oncogene signaling supported by two surface antigens and also possesses the ability to stimulate the immune system through its Fc domain, could be a superior combination therapy and provide synergistic effects in a single molecule. Here, we initially tested this hypothesis by generating different monospecific antibodies (anti-EGFR, anti-HER2) and bispecific antibodies (anti-EGFR/HER2) using knob-into hole and CrossMab technology, completed the functionality and characterization of the engineered protein, proved that FcR interactions contribute to the stimulatory effect, and finally observed the synergistic therapeutic effect *in vitro*.

## Results and Discussion

### Construction and Purification of Bispecific Antibodies

The pcGS vector backbone had been used to clone monospecific and bispecific antibodies to check protein expression. In this study, we successfully designed and expressed a bispecific antibody targeting EGFR and HER2 using a single-vector strategy that incorporates the knobs-into-holes (KiH) heterodimerization system. Specific engineered alterations (S354C/T366W for the “knob” and Y349C/T366S/L368A/Y407V for the “hole”) were introduced into the CH3 domains to promote selective heavy chain heterodimerization, consistent with prior studies validating the effectiveness of the KiH approach.^27,28^ Western blot analysis confirmed the expression of intact bispecific IgG molecules at approximately 180 kDa in CHO cells, indicating correct assembly. Notably, the comparable intact mass between the monospecific and bispecific antibodies suggests that the introduced mutations did not adversely affect overall molecular weight or structural integrity. However, the partial fragmentation observed in the bispecific antibody may reflect reduced molecular stability, a phenomenon previously associated with engineered IgGs that involve modifications to the CH3 region.^29^ Importantly, the single-vector system facilitated the co-expression of all four polypeptide chains within a single construct, simplifying production relative to conventional co-transfection strategies. This streamlined approach reduces variability related to transfection efficiency and plasmid copy number. Gene organization within the plasmid is depicted in Figures 1A and 1B. Western blot results (Figure 1C) confirmed bispecific IgG expression, as indicated by a band above 180 kDa. These data support the feasibility of producing bispecific antibodies using engineered CH3 interfaces in a single-vector format within CHO cells.

**Figure 1.**
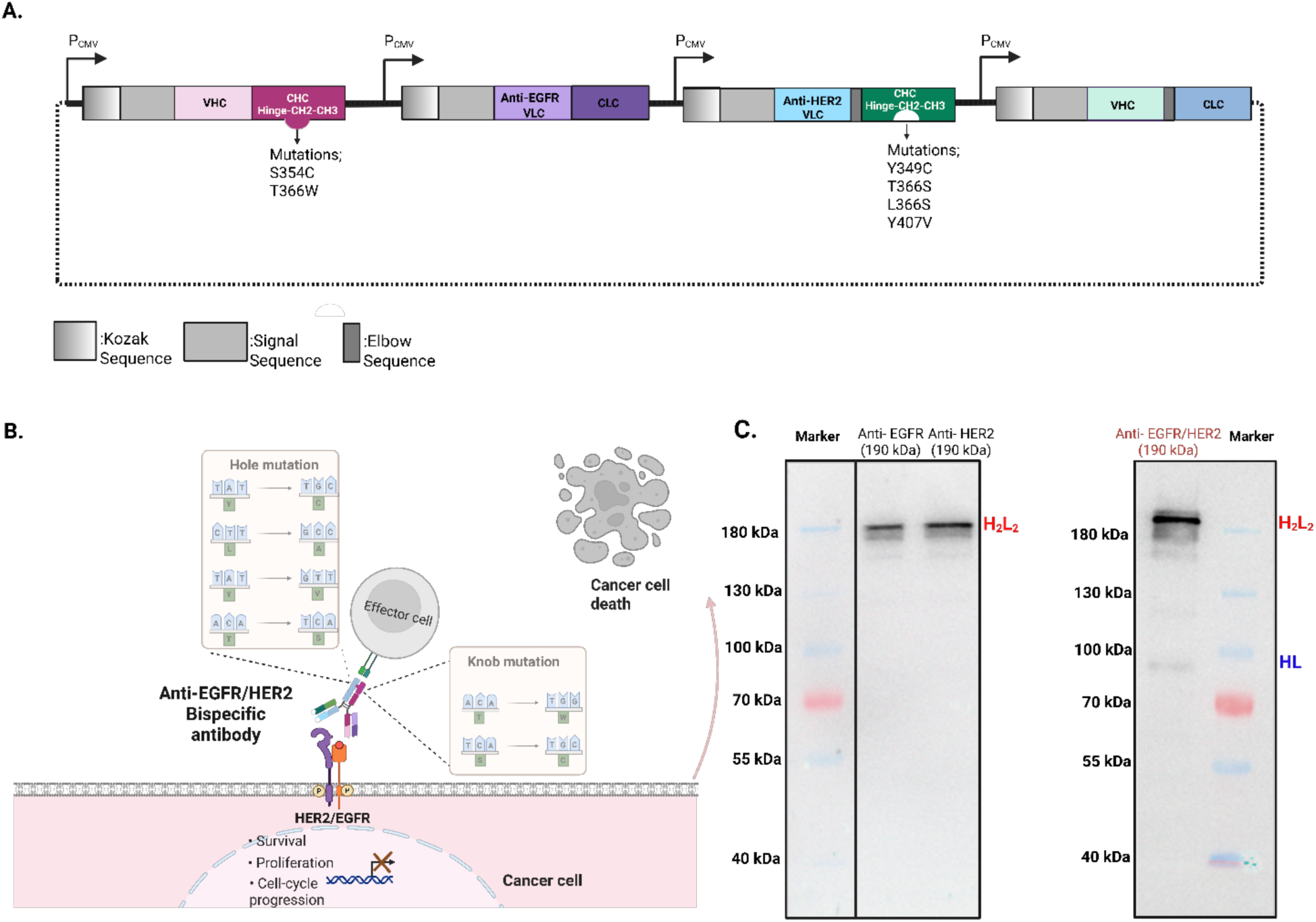
Expression of recombinant monospecific and bispecific antibodies. (A) Orientation and construction of recombinant monospecific antibodies, expressed in CHO cells. (B) Orientation and construction of recombinant bispecific antibodies with engineered arms (Created with BioRender.com). (C) Western blot analysis of a stable pool sample of monospecific anti-EGFR, anti-HER2, and bispecific anti-EGFR/HER2. These samples were collected from a stable pool. Western blot analysis for monospecific antibodies and the bispecific antibody (kDa as molecular size). All antibodies were observed above 180 kDa. CHC, VLC, CLC, and VHC represent constant heavy chain, variable light chain, constant light chain, and variable heavy chain, respectively. From top to bottom, the red arrows represent intact antibody mass. Whereas the H2L2 label indicates a complete antibody, the HL label represents a half-antibody species consisting of only one heavy and one light chain.

The primary aim of ELISA was to evaluate the binding activities of the developed monospecific and bispecific antibodies to their target antigens. The monospecific antibodies demonstrated expected target binding using commercially available kits, which rely on Fc-specific HRP-conjugated secondary detection. These results validated the correct folding and antigen recognition ability of the engineered variable domains, utilizing recombinant IgG molecules to confirm antigen specificity. Commercial ELISA kits were used to determine the binding affinity of monospecific antibodies, thereby evaluating the binding capacity of these antibodies to their target antigens. To overcome the inherent limitations of conventional ELISA formats in detecting bispecific antibody binding, particularly due to the reliance on Fc recognition, we performed a sandwich ELISA platform. This approach enabled us to confirm the simultaneous engagement of both HER2 and EGFR antigens, a functional hallmark of bispecific antibodies (Figure 2A-C). The successful detection of a signal in this format suggests that both antigen-binding arms retained their specificity and were functionally accessible, even within the context of a heterodimeric IgG molecule.

**Figure 2.**
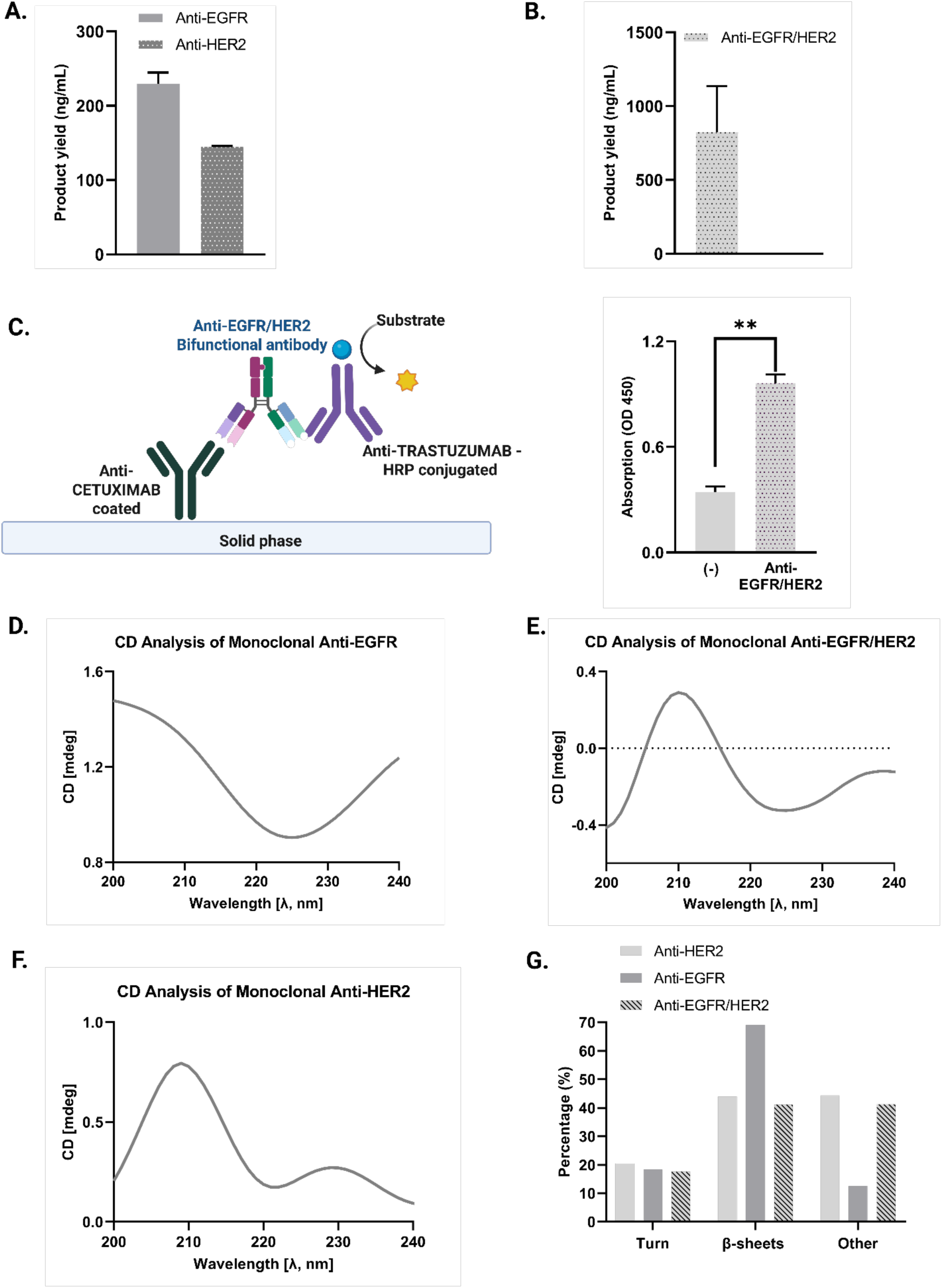
Monospecific antibodies targeting Anti-EGFR and Anti-HER2, along with the bispecific antibody targeting Anti-EGFR/HER2, were evaluated for their specific binding affinities to the corresponding target antigens. Absorbance values were measured at 450 nm for each sample, and binding affinities were calculated by normalizing the obtained data to their respective negative control groups in each experimental set. n = 2 replicate means are presented. Statistical analysis was conducted using a t-test on ELISA measurements to ascertain significant variances from negative control values. Statistical significance was established accordingly. (A) Monospecific antibodies were quantified using ELISA kits, with measurements obtained after the standard curve was constructed. (B) The yield of bispecific antibody Anti-EGFR/HER2 was quantified using a commercial CETUXIMAB ELISA kit. (C) A schematic representation was provided, followed by the determination of the absorbance values at 450 nm for the Anti-EGFR/HER2 antibody. These values for the bispecific antibody, which targets EGFR, showed a significant increase compared to the negative control (**p < 0.01). (D-F) CD spectral data of purified native monospecific and bispecific antibodies. (G) The graphic of fold recognition results was analyzed using the online tool BeStSel.^33^

Circular dichroism (CD), a widely used method for studying the secondary structures of proteins, was employed to evaluate changes in the secondary structural conformation of the produced native monospecific antibodies and the bispecific antibody. A wavelength range from 200 to 240 nm was used to determine secondary structures. Evaluating these spectral features played a crucial role in determining whether the interface mutations compromised the structural integrity of the protein. Our results showed that the CD spectrum of the Anti-EGFR/HER2 bispecific antibody maintained the typical secondary structure of a β-sheet protein, verifying that the introduced engineered parts had no significant impact on its structural conformation. The spectra appeared to be quite similar, exhibiting features characteristic of proteins, with a negative band at approximately 220 nm and a positive band at around 208 nm (Figure 2D-F). The observation that the anti-EGFR antibody exhibits a spectral peak near 200 nm aligns with previously reported CD profiles of Cetuximab, further supporting the expected β-sheet–dominated secondary structure.^30^ The BestSel online tool was used to predict secondary structures. The results also confirmed that % antiparallel β-sheets and β-turns structure predominated in all developed antibody structures (Figure 2G). Therefore, it has been demonstrated that engineered parts do not cause significant changes in the secondary structural conformation, and it has been confirmed that the protein remains structurally and conformationally intact. Similar findings have been reported in other antibody constructs, where conformational stability was preserved despite mutations within the CH3 interface.^31,32^

### Immunocytochemistry revealed distinct antibody–receptor binding patterns across the three cell lines

The binding specificity of the engineered antibodies toward their cognate receptors was assessed using an ICC assay (Figure 3). Because CHOK1 cells were used as a production host to express the antibodies, this cell line was also used for ICC testing (Figure S6). CHOK1 cells are intrinsically devoid of endogenous EGFR and HER2 expression; thus, no signal related to the presence of either receptor was expected when incubating with anti-EGFR, anti-HER2, or bispecific anti-EGFR/HER2 antibodies. Immunodetection of β-actin served as a positive control for protein expression within cells and to confirm that the ICC workflow was appropriate. Nuclear counterstaining with DAPI was conducted under all conditions to ensure the presence and viability of cells. Confocal microscopy analysis revealed strong DAPI staining signals across the samples, indicating successful nuclear labeling. The detection of β-actin yielded a strong FITC signal from the secondary antibody, validating the assay’s performance. In contrast, ICC with anti-EGFR, anti-HER2, and bispecific antibodies resulted in undetectable levels of FITC signal, yet DAPI staining remained intact. The absence of fluorescence is consistent with the known receptor-negative status of CHOK1 cells and confirms the lack of off-target or nonspecific binding of the tested antibodies (Figure 3A).

**Figure 3.**
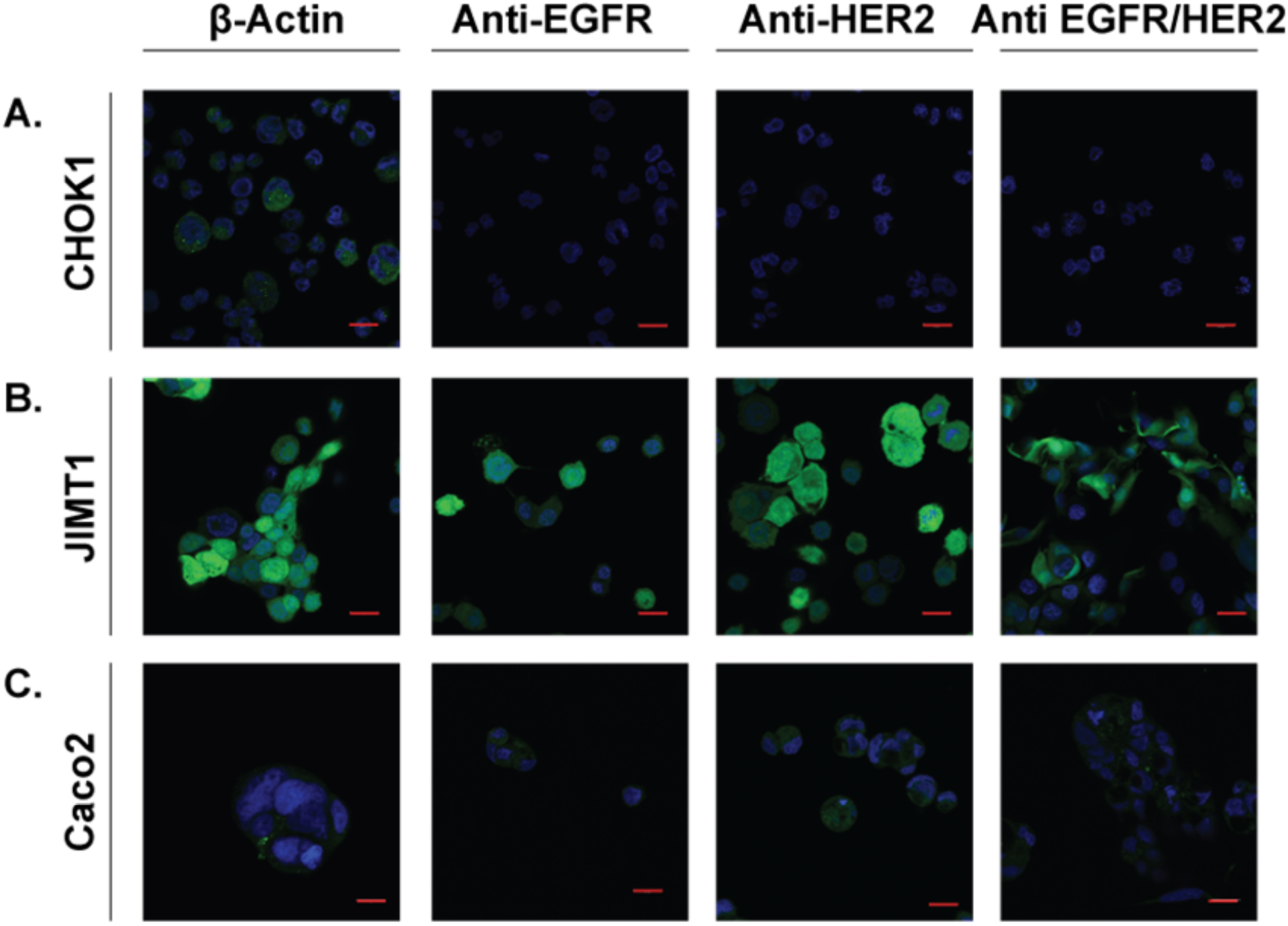
High-resolution immunocytochemistry analysis of antibody–receptor interactions in three cell lines with distinct EGFR/HER2 expression profiles (A) CHOK1 cells, which lack endogenous EGFR and HER2 expression, show only DAPI nuclear staining (blue) and no detectable FITC signal (green) following incubation with anti-EGFR, anti-HER2, or bispecific anti-EGFR/HER2 antibodies, confirming the absence of receptor-dependent membrane binding. (B) JIMT-1 cells, characterized by high EGFR and moderate HER2 expression, exhibit strong and membrane-localized FITC fluorescence, consistent with robust binding of anti-EGFR and bispecific antibodies, and moderate binding of anti-HER2. These images highlight the expected receptor abundance hierarchy and validate antibody specificity under high-magnification confocal imaging. (C) Caco-2 cells, which form tightly packed polarized epithelial aggregates, display markedly reduced FITC fluorescence across all antibody conditions. This diminished signal is consistent with restricted antibody penetration and limited apical receptor accessibility characteristic of differentiated Caco-2 cultures. DAPI staining is visible across all panels, confirming the integrity of the nuclear structure. All images were acquired using a 63× oil-immersion objective, with scale bars representing 20 µm in length.

Taken together, these findings confirm that CHOK1 cells represent a suitable expression platform for producing antibodies against either EGFR or HER2, as the absence of endogenous receptors avoids unwanted antibody–receptor interactions that could otherwise compromise expression efficiency or confound downstream analyses.

Immunocytochemistry was conducted using the JIMT-1 breast cancer cell line, which endogenously expresses both EGFR and HER2, to evaluate receptor-dependent binding of the engineered antibodies (Figure S7). Based on this receptor profile, anti-EGFR, anti-HER2, and bispecific anti-EGFR/HER2 antibodies were expected to show a strong cell-surface binding. These expectations were confirmed by confocal imaging. In all conditions of treatment with antibodies, DAPI staining of cell nuclei was distinct, thereby ensuring sample integrity for the reliable assessment of receptor-associated fluorescence. FITC signals, reflecting antibody binding, were strongly present at the cell membrane. Treatment with the bispecific anti-EGFR/HER2 antibody resulted in substantial and widespread fluorescence, indicating its ability to bind to both receptors simultaneously. Observed fluorescence intensities follow the known receptor distribution, further supporting the biological relevance and binding fidelity of the antibody panel (Figure 3B).

For further characterization of receptor accessibility and antibody–receptor interaction, ICC was performed using the Caco-2 colorectal adenocarcinoma cell line (Figure S8). However, unlike CHOK1 or JIMT-1 cells, Caco-2 cells exhibit a highly polarized, tightly packed epithelial morphology and spontaneously form dense three-dimensional aggregates and dome-like structures under standard culture conditions. This architectural organization is well-documented to hinder the diffusion of antibodies and reduce the effective exposure of cell-surface receptors, particularly in ICC or live-cell labeling workflows.^34^

Indeed, consistent with these reports, confocal imaging demonstrated that DAPI staining successfully highlighted cell nuclei across all samples. In contrast, FITC-associated membrane staining was relatively less pronounced for each of the three antibodies, including anti-EGFR, anti-HER2, and the bispecific anti-EGFR/HER2 antibody. Despite these structural limitations, mild fluorescence patterns corresponding to anti-EGFR and bispecific antibody binding were detectable in less compact regions of the culture, consistent with the presence of EGFR on accessible membrane surfaces. Together, these findings support the conclusion that the diminished FITC signal is primarily due to the epithelial organization and aggregation state of Caco-2 cultures rather than the absence of receptor expression (Figure 3C). This observation is consistent with the prior literature, which describes restricted antibody accessibility in polarized intestinal epithelial models.

### Dual Targeting Decreases Cell Viability in EGFR/HER2-Expressing Cell Lines

The cytotoxic effects of anti-EGFR, anti-HER2, and bispecific anti-EGFR/HER2 antibody treatments were evaluated by MTT assay across multiple cell lines, with CHOK1 cells serving as a negative control. As expected, CHOK1 cells (Figure 4A) displayed no significant changes in viability following treatment with any of the antibody formats at 1 µg/mL or 10 µg/mL, confirming the target specificity of the antibodies. The lack of cytotoxicity in this non-tumorigenic line, which does not overexpress EGFR or HER2, supports the selective nature of the observed therapeutic effects. This proves that specific binding to the antigen has occurred.

**Figure 4.**
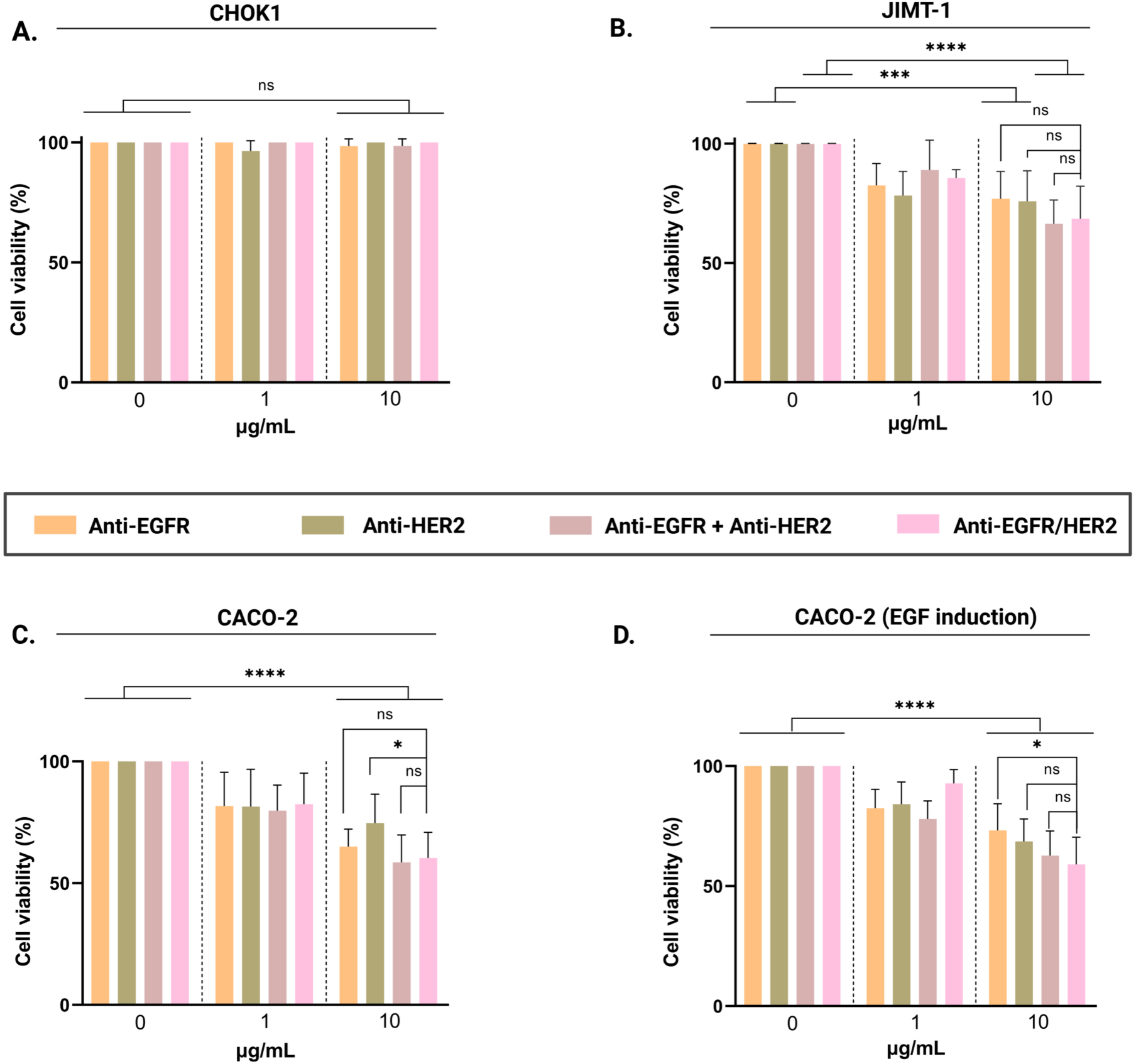
The *in vitro* antitumor activity of the Anti-EGFR/HER2 bispecific antibody was evaluated by various assays. The effects of monoclonal antibodies on the proliferation of Caco-2 cells stimulated by EGF were assessed, with experiments performed in the absence or presence of EGF. (A) CHOK1 (B) JIMT-1 (C) Caco-2 cells (uninduced) and (D) Caco-2 cells induced with EGF were treated with different antibody concentrations (1 µg/ml and 10 µg/ml). All monospecific and bispecific antibodies were added for 24 h before the MTT assay was performed to determine cell viability compared to the control group. In the experiments, the combined treatments of monospecific antibodies designated as Anti-EGFR+Anti-HER2. ***p<0.001, **p<0.01, *p<0.05, and ns: no significant difference. The data obtained as a result of the experiments performed with three independent biological replicates.

In contrast, JIMT-1 cells (Figure 4B) showed a significant reduction in cell viability upon exposure to antibody treatments. Monotherapies with anti-EGFR and anti-HER2 led to a moderate loss of viability at a concentration of 10 µg/mL. In contrast, combination therapy and the bispecific anti-EGFR/HER2 antibody produced the most pronounced cytotoxic responses (p < 0.001 and p < 0.0001, respectively). This observation supports the concept that dual receptor blockade can effectively bypass acquired resistance mechanisms.^20,35–37^

Although effects at 1 µg/mL were less prominent, they followed a similar trend. Interestingly, the cytotoxic outcomes for the bispecific and combination treatments were statistically nonsignificant, suggesting comparable efficacy between these dual-targeting strategies. If HER2 maintains autonomous signaling activity independent of EGFR, particularly in resistant cell lines such as JIMT-1^38^, then the suppressive impact of the bispecific antibody may be limited, despite the presence of EGFR-HER2 interactions. The lack of significant difference among the four treatment modalities, despite some decrease in viability, may reflect the limited capacity of the bispecific antibody to redirect signaling to HER2 via EGFR in the context of HER2 dominance.

Under basal conditions, Caco-2 cells (Figure 4C) responded similarly. All therapies induced significant reductions in viability at 10 µg/mL (p<0.0001). Greater effects were observed with dual antibody approaches; however, no statistically significant difference was detected between combination and bispecific formats. These findings indicate that dual targeting confers enhanced cytotoxicity compared to single-agent therapies, although the extent of this enhancement may vary depending on the antibody configuration. The bispecific antibody induced a significantly greater reduction in cell viability compared to anti-HER2 monotherapy (p<0.05). However, a similar effect was not observed when comparing anti-EGFR treatment with the bispecific antibody. While EGFR expression is high in Caco-2 cells, HER2 expression is generally limited.^39^ Therefore, it is logical that the difference is seen through HER2.

Upon EGF stimulation (Figure 4D), Caco-2 cells exhibited a partial resistance to antibody-induced cytotoxicity, consistent with prior observations of EGF-mediated pro-survival signaling.^40^ Despite this, anti-EGFR alone and both dual-targeting approaches still produced significant reductions in viability relative to controls (p<0.0001). At 10 µg/mL, combination and bispecific antibody treatments remained superior to monotherapies, though again without a statistically meaningful difference between the two. Collectively, these results demonstrate that antibody-mediated cytotoxicity is both target- and dose-dependent, and that dual receptor blockade provides a distinct advantage over monotherapies. At 10 µg/mL, in ligand-dependent cultured Caco-2 cells, bispecific antibody treatment produced a statistically significant difference (p<0.05) only when compared to anti-EGFR monotherapy. Notably, HER2’s heterodimerization with EGFR prolongs proliferative signaling by preventing ligand-bound EGFR from undergoing endocytosis.^40^ This interaction increases the effect of EGF, increasing the growth sensitivity of cells with high HER2 expression. When considered in Caco-2 cells, dual-targeting approaches maintained their superiority over various monotherapies in both basal and ligand-stimulated conditions. It has been noted that EGF causes an increase in the heteroassociation of EGFR and HER2 and the homoassociation of HER2 within minutes, and that EGF stimulation also increases the size of HER2 clusters.^41^ This phenomenon is closely related to the receptor density of the cell lines. Different expression levels of HER2 and EGFR lead to therapeutic response heterogeneity, supporting the development of resistance to monospecific agents.^37,42^ The literature explains that EGF stimulation effectively induces homodimerization of EGFR.^40,43,44^ However, there are two different findings on how EGF affects HER2. While it is stated that EGF binds strongly to EGFR and activates it, leading to the formation of EGFR/HER2 heterodimers,^44^ there is another literature showing that EGF cannot trigger the formation of the EGFR/HER2 complex and that the interaction between EGFR and HER2 is much weaker than that of the EGFR homodimer.^40^ Based on our findings, EGF appears to have functionally engaged HER2 in the signaling process, making the observed significant difference in response to anti-EGFR treatment a predictable outcome.

### Antibody-Mediated Inhibition of Metabolic Activity in 3D Tumor Cell Spheroids

Three-dimensional spheroids were successfully generated in both the experimental and control cell lines (Figure 5A). Following spheroid formation, metabolic activity was quantified using the CellTiter-Glo 3D cell viability assay, which measures ATP-dependent luminescence as an indicator of viable cells. The resulting viability profiles are presented in Figure 5B. For the JIMT-1 cell line, fold-change analysis revealed a progressive and consistent reduction in spheroid viability across all antibody-treated groups, with the most pronounced decrease observed in the antibody-treated spheroids. As expected, JIMT-1 cells exhibited sensitivity to all antibody conditions, with each demonstrating a statistically significant reduction in viability compared to the negative control (p < 0.0001). Notably, only 10 µg/mL anti-EGFR/HER2-treated spheroids showed a significantly greater decrease in viability compared with the 10 µg/mL anti-HER2-treated group (p<0.05). This pattern aligns with our immunocytochemistry findings, suggesting that the enhanced effect is driven by EGFR abundance and the resulting synergistic contribution of dual-target engagement.

**Figure 5.**
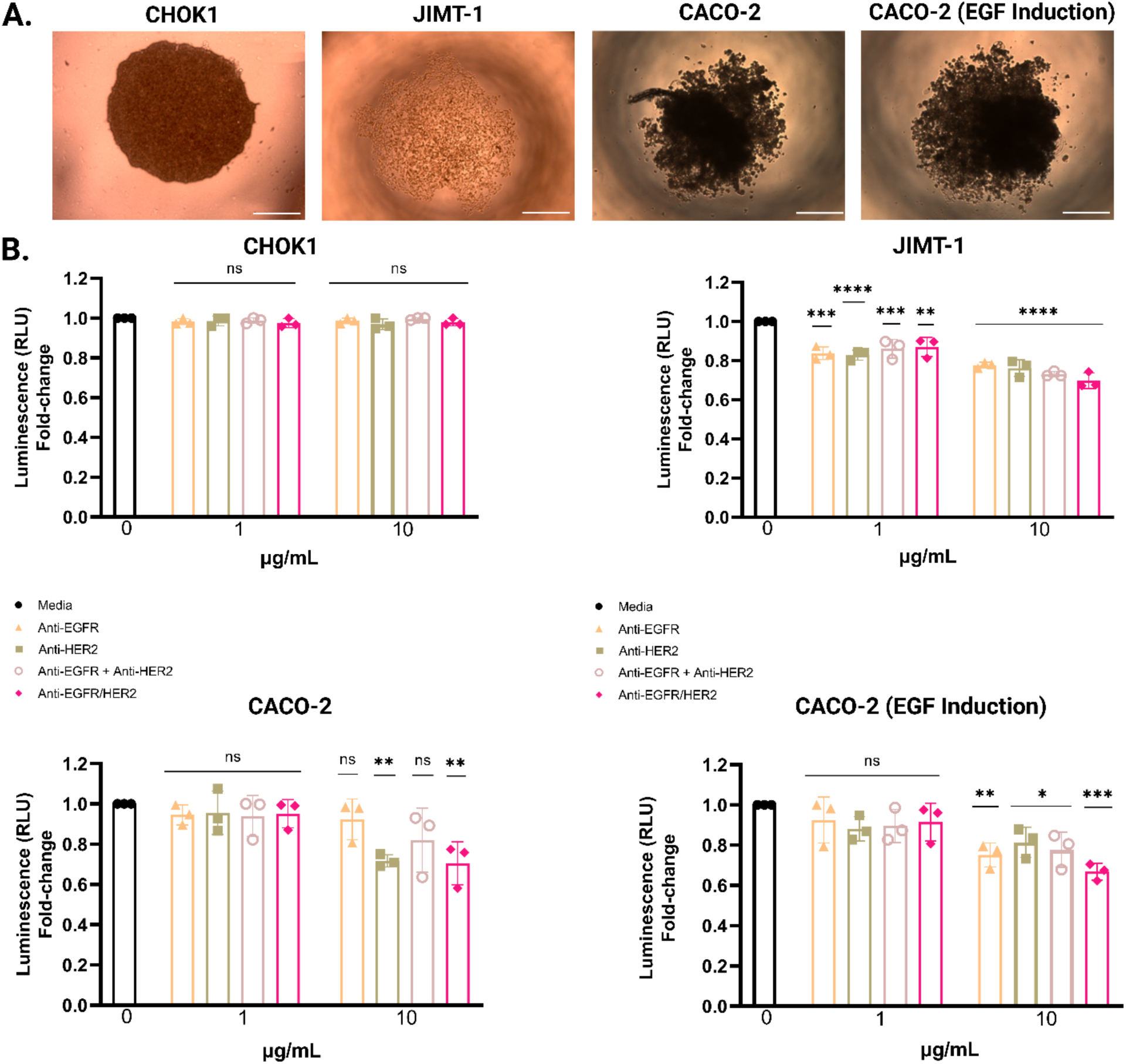
3D cell viability assessment was performed using the CellTiter-Glo 3D assay on CHOK1, Caco-2 and JIMT-1 spheroids. (A) Confocal microscopy images obtained following 24 hours of spheroid culture (scale bar = 500 µm). (B) One-way ANOVA was applied using the no-antibody media condition as the control group for Caco-2, Caco-2 treated with EGF, and JIMT-1 spheroids. Statistical significance is indicated as ****p<0.0001, ***p<0.001, **p<0.01, *p<0.05, and ns: no significant difference. Data represent the mean ± SD from experiments performed in three replicates.

In the Caco-2 spheroids, the viability pattern differed depending on EGF stimulation. In the absence of EGF, Caco-2 cells unexpectedly displayed higher responsiveness to HER2-targeting treatment. However, the response shifted to the expected pattern upon EGF stimulation. The bispecific antibody induced a significantly greater reduction in viability (p<0.001) compared with anti-EGFR alone, with 10 µg/mL (p<0.01), and both treatments produced more substantial effects relative to the negative control. These results suggest that ligand-mediated receptor activation enhances EGFR accessibility, leading to a more pronounced therapeutic response.

### Bax/Bcl-2 Ratio Shifts Indicate Enhanced Apoptosis Following Antibody Treatments

To evaluate the apoptotic response following antibody treatment, we measured the Bax/Bcl-2 mRNA expression ratios as an indicator of apoptosis induction in JIMT-1 and Caco-2 cell lines (Figure 6A–C). In JIMT-1 cells (Figure 6A), treatment with anti-EGFR/HER2 bispecific antibody resulted in a higher apoptotic response, with a significant difference (p<0.0001) compared to untreated controls. Notably, the combination treatment (anti-EGFR/HER2) exhibited a stronger apoptotic response than anti-EGFR monotherapy alone (p<0.0001), suggesting a potential synergistic effect of dual targeting in JIMT-1 cells. In Caco-2 cells (Figure 6B), the bispecific anti-EGFR/HER2 treatment significantly elevated apoptotic gene expression compared to untreated controls (p<0.001), as well as compared to monotherapies (anti-EGFR alone, p<0.05). In Caco-2 cells stimulated with EGF (Figure 6C), the apoptotic response was less pronounced. Treatment with anti-EGFR and anti-HER2 individually or in combination moderately increased Bax/Bcl-2 ratios compared to untreated control cells. However, significant differences were observed only when comparing the bispecific treatment group with untreated controls (p < 0.01). These findings again indicate an enhanced apoptotic effect mediated by simultaneous targeting of both EGFR and HER2 receptors in Caco-2 cells. Cropped gel images are shown for comparison between groups (Figure S1A–C).

**Figure 6.**
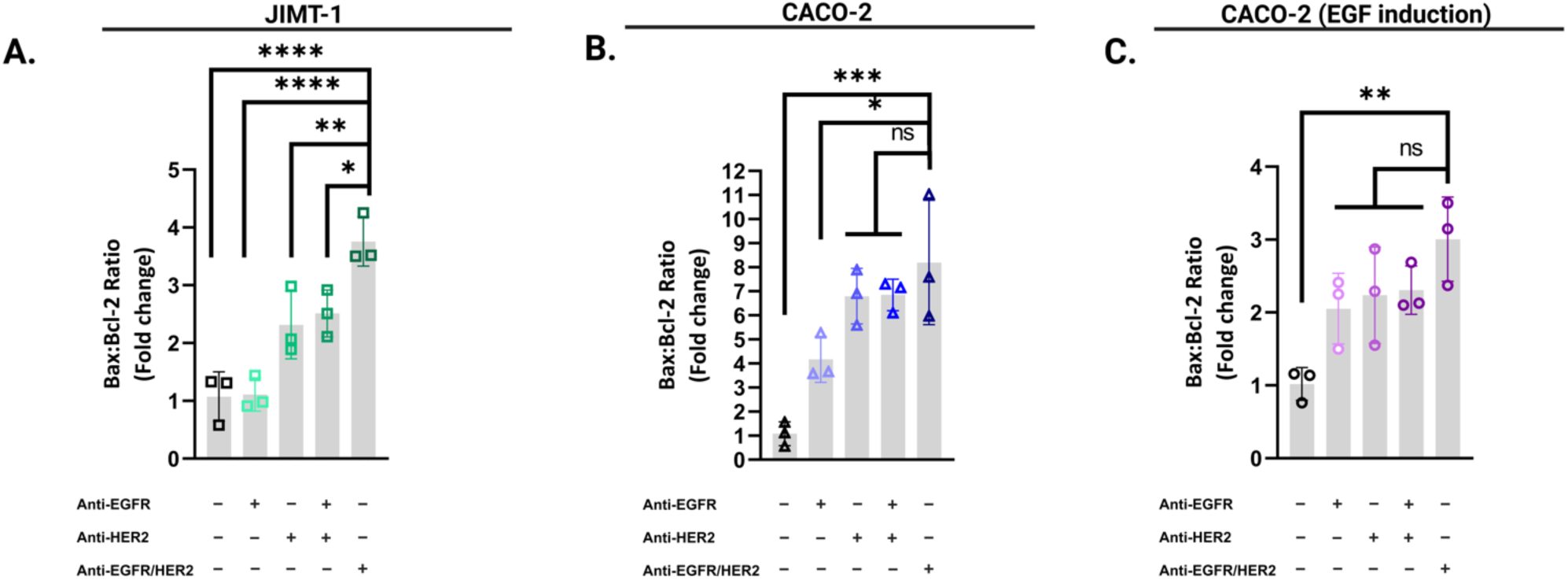
Bax/Bcl-2 ratio analysis reveals enhanced apoptotic signaling. (A-C) Impact of Anti-EGFR, Anti-HER2, and Anti-EGFR/HER2 antibodies on Bax and Bcl-2 mRNA expression levels in Caco-2 and JIMT-1 cells. Cells were incubated with antibodies for 18 hours. The mRNA expression was quantified via qRT-PCR and normalized against Actin. Dots depict technical replicates, while bars indicate mean ± SD. Data are presented as fold change relative to untreated controls with mean ± S.E. (n = 3). Statistical analysis for multiple comparisons was conducted using one-way ANOVA.

These findings are consistent with studies showing that Bcl-2 downregulation and Bax activation promote mitochondrial apoptosis.^45,46^ Notably, the Bcl-2 protein exhibits minimal expression in JIMT-1 cells,^47^ a finding that aligns well with our own experimental results. In trastuzumab-resistant HER2-positive xenografts, altering the Bax/Bcl-2 balance has been shown to restore apoptosis, thereby improving mitochondrial apoptotic signaling and leading to tumor regression.^48^ Our data support this mechanism in JIMT-1 cells, where upregulation of Bax relative to Bcl-2 may be a key determinant of treatment response. In contrast, Caco-2 cells exhibited moderate increases in Bax/Bcl-2, which were significant only under bispecific treatment (p < 0.001), indicating that dual blockade amplifies intrinsic apoptosis even in EGFR/HER2 co-expressing models. However, our observation that EGF induction in Caco-2 cells attenuates the apoptotic response to antibody treatment is noteworthy, reflecting literature that highlights the adaptive responses of cancer cells in ligand-rich microenvironments. The involvement of EGF in modulating apoptotic signaling pathways is well established. EGF exposure has been reported to promote cellular proliferation, suppress apoptosis, and downregulate Bax expression, while concurrently inducing anti-apoptotic proteins.^49^ Moreover, apoptosis triggered by EGF has been shown to proceed through caspase-independent mechanisms, characterized by conformational activation, oligomerization, and mitochondrial translocation of Bax in cells with elevated EGFR expression.^50,51^

These observations are in agreement with our MTT results, where reduced cell viability, measured as decreased metabolic activity, correlates with apoptotic induction. The observed differences between MTT assay outcomes and apoptotic marker expression likely reflect variations in the timing and mechanisms of cell death. MTT assays indicate mitochondrial activity, which may remain detectable during early apoptotic stages, potentially resulting in a slight overestimation of viable cells. Moreover, cell-to-cell variability, particularly under EGF stimulation, may influence these measurements and contribute to variability in statistical significance. These observations are consistent with prior reports suggesting that apoptotic signaling can precede detectable metabolic decline, highlighting the multifactorial and dynamic nature of cell death processes.^52^

### Antibody-Triggered Cytotoxic Response Mediated by CD14⁺ Monocytes

Calcein release assays were utilized to determine whether antibody-induced cytotoxicity involved Fc receptor-mediated interactions, with CD14⁺ monocytes serving as effector cells in co-culture with fluorescently labeled target cells (Figure 7A). In JIMT-1 cells, treatment with anti-EGFR, anti-HER2, or combined antibody approaches resulted in a significant increase in cytotoxicity compared to untreated controls (**p<0.01) (Figure 7B). Notably, cytotoxicity levels were similar across all antibody formats, suggesting effective engagement of monocytes through Fc–Fcγ receptor interactions. Similarly, in Caco-2 cells under basal conditions, antibody treatments induced significant cytotoxicity compared to untreated cells (**p<0.01; ***p<0.001) (Figure 7C). Combined and bispecific antibody treatments elicited slightly higher cytotoxic responses than monotherapies, although differences among antibody formats were not statistically significant. In EGF-stimulated Caco-2 cells, antibody-mediated cytotoxicity remained significant compared to controls (**p<0.01) (Figure 7D). However, overall cytotoxicity levels appeared slightly elevated compared to basal conditions, indicating that ligand activation may sensitize cells to Fc-mediated killing, potentially due to receptor upregulation. These results confirm that antibody Fc domains effectively engage CD14⁺ monocytes, promoting cytotoxic responses against EGFR/HER2-expressing tumor cells. The ability of therapeutic antibodies to mediate cytotoxicity through immune effector cells, such as monocytes, is a crucial mechanism that complements their direct tumoricidal effects. In the present study, calcein release assays demonstrated that anti-EGFR, anti-HER2, and bispecific anti-EGFR/HER2 antibodies effectively engaged CD14⁺ monocytes to mediate Fc-dependent cytotoxicity against tumor cells.

**Figure 7.**
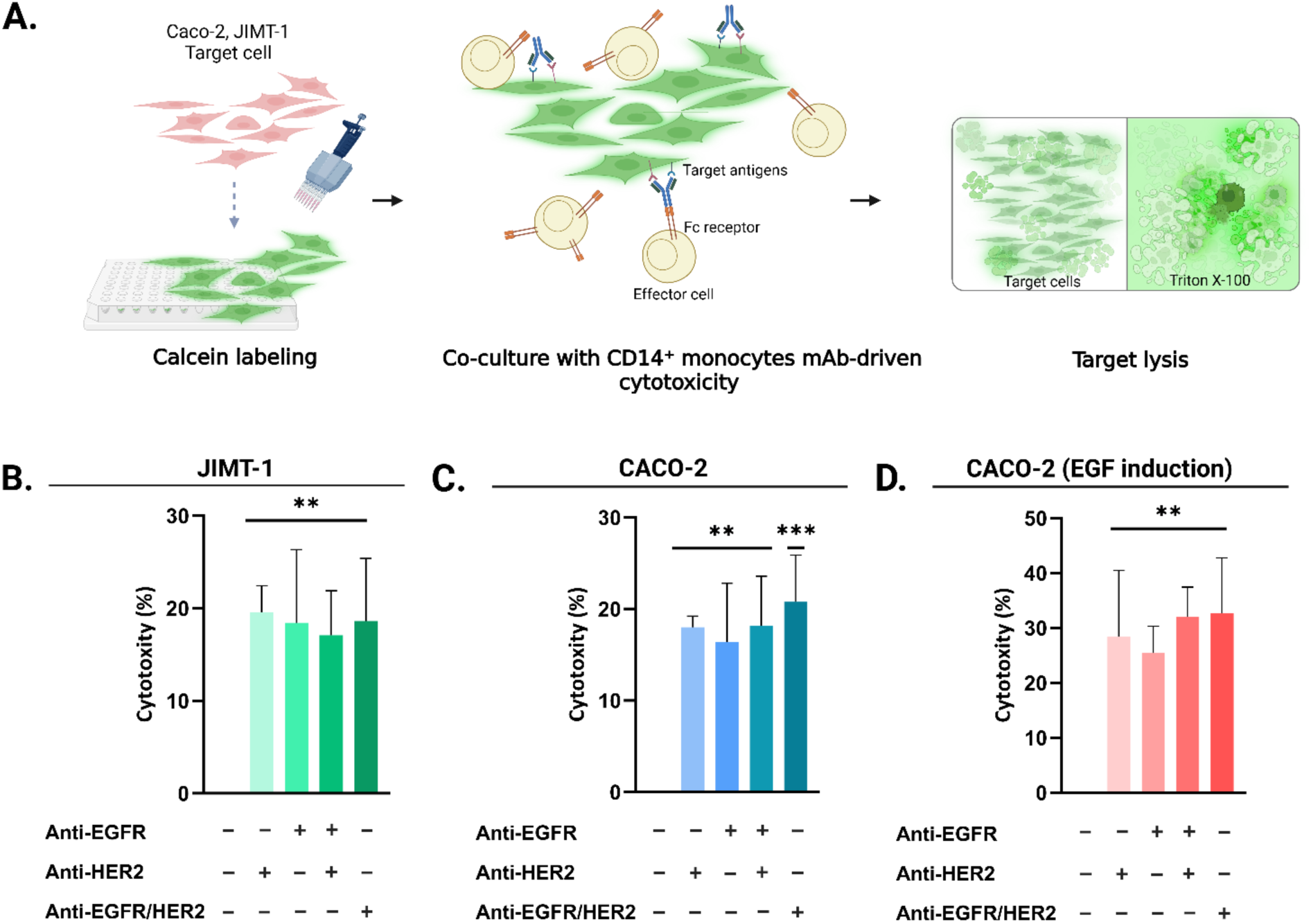
Schematic representation and quantitative results of calcein release assay evaluating antibody Fc-mediated cytotoxicity. (A) Labeled target cells were co-cultured with CD14⁺ monocytes to evaluate antibody Fc-mediated cytotoxicity by quantifying calcein release into the extracellular medium upon loss of membrane integrity. Monocytes were engaged via the Fc regions of therapeutic antibodies targeting EGFR and HER2. (B-D) Antibody binding and Fc engagement led to immune-mediated cytotoxicity, resulting in the release of calcein into the culture medium. Graphs show the percentage of cytotoxicity in JIMT-1, Caco-2, and EGF-stimulated Caco-2 cells following treatment with anti-EGFR, anti-HER2, combined antibodies, and bispecific anti-EGFR/HER2 antibodies. Data are presented as mean ± SD (n = 3). Statistical significance was evaluated using one-way ANOVA with multiple comparisons (**p<0.01; ***p<0.001; ns: not significant).

Previous studies have highlighted the role of Fc-FcγR interactions and target antigen density as critical determinants of monocyte-mediated ADCC and ADCP.^53–55^ While CD14⁺ monocytes are recognized for their capacity to detect and phagocytose opsonized tumor cells,^56^ their ability to mediate ADCC has remained controversial. For instance, Trivedi et al. (2016)^57^ reported limited cytotoxic activity by CD14⁺ monocytes in the presence of EGFR-targeting antibodies. Nevertheless, our results indicate that CD14⁺ monocytes can indeed participate in effective antibody-dependent cytotoxicity when engaged by anti-EGFR, anti-HER2, or bispecific antibodies. These findings suggest that, under appropriate conditions, such as sufficient target antigen density and optimized antibody configuration, CD14⁺ monocytes can exert Fc-mediated effector functions, thereby contributing significantly to the cytotoxic response.

## Conclusion

We produced an anti-EGFR/HER2 bifunctional antibody in suspension CHO cells by forming heterodimers using knob into hole technology. The bifunctional antibody, evaluated for its synergistic effect of dual targeting, was tested in cell lines exhibiting receptor heterogeneity or resistance to single-agent therapies. Our data suggest that bispecific antibody therapy targeting EGFR and HER2 provides potent antitumor effects through two mechanisms: induction of apoptosis and recruitment of immune effectors. Dual EGFR/HER2 blockade via bispecific antibodies represents a promising strategy to overcome ligand-induced resistance, restore apoptotic signaling, and facilitate immune-mediated killing in solid tumors that express these receptors.

## Materials and Methods

### Materials

Cell strains were specified as *E. coli* DH5α, Chinese hamster ovary (CHO) cells, and Caco-2, JIMT-1, and CHOK-1, THP1-Dual™ MD2-CD14-KO-TLR4 Cells (Invivogen) for cloning, protein expression studies, and *in vitro* cell assays, respectively. Caco-2 was kindly provided by Sreeparna Banerjee from Middle East Technical University, Ankara, Turkey. mAb-Based ELISA Kit (*ImmunoGuide* Trastuzumab ELISA, IG-AB105), Cetuximab mAb-Based ELISA Kit *(ImmunoGuide* Cetuximab ELISA, IG-AB112), were purchased for ELISA. Horseradish peroxidase (HRP) conjugate anti-Trastuzumab (10 µg/mL, 30 mL volume) and anti-Cetuximab mouse monoclonal antibody (600 µg per vial), both provided as gifts by AybayTech Biotechnology, were used for dual-binding ELISA. For western blotting, goat pAb Anti-Human IgG (HRP) (ab98624, Abcam) was preferred. For purification, HiPrep 16/60 Sephacryl S-200 HR column (Cytiva) and 1mL Cytiva HiScreen Capto L HiTrap Protein L were used. For immunocytochemistry, collagen (C8919, Sigma-Aldrich), formaldehyde (923.012, ISOLAB), Triton X-100 (T8787, Sigma Aldrich), β-Actin (8H10D10) Mouse mAb (3700, Cell Signaling), Goat anti-Mouse IgG (H+L) Secondary Antibody, FITC (31569, Thermo Fisher Scientific), Alexa Fluor® 488 anti-human IgG Fc Antibody (410705, Biolegend), Ibidi Mounting Medium With DAPI (50011, Ibidi), anti-EGFR (1.8 mg/ml), anti-HER2 (1.8 mg/ml), and anti-EGFR/HER2 (1.8 mg/ml) were used. For 3D cell culture, CellTiter-Glo® 3D Cell Viability Assay (Promega) was utilized.

## Methods

### Construction of DNA Sequences Encoding Bispecific Antibody

Gene fragments for both engineered and wild-type forms of antibody heavy and light chains were synthesized by Genewiz (Genewiz, NJ) and inserted into the pcGS plasmid backbone via Gibson assembly. Overhang regions were incorporated into gene fragments to facilitate this process. All plasmid and primer designs were conducted *in silico* using Benchling. For the native anti-HER2 plasmid, the vector backbone was amplified in two segments to include separate promoter, enhancer, terminator, and selection marker elements using BB-PCR1-F/BB-PCR1-R and BB-PCR2-F/BB-PCR2-R primer pairs. The light chain gene was amplified using NLF and NLR primers with overhangs compatible with BB-PCR2-F and BB-PCR1-R. The heavy chain was amplified using NHF, NHR, Fcc-F, and Fcc-R primers, and the fragments were fused using PCR Assembly to obtain the native Fc region. These four components, two backbone regions, heavy, and light chains- were assembled into the final plasmid using Gibson Assembly. Separate signal sequences for both chains and Kozak sequences were added to the N-termini to enhance expression and secretion. For bispecific antibody cloning, Anti-HER2-CrossMab-Hole and Anti-EGFR-Knob plasmids were first constructed individually. To create a single bispecific plasmid, the Anti-EGFR-Knob construct was linearized with PacI. Two fragments were PCR-amplified from the Anti-HER2-CrossMab-Hole plasmid: the HER2-binding domain (PCR1, using SP-F and SP-R) and an additional CMV promoter region (PCR2, using SP-PCR2-F and SP-PCR2-R). These, along with the linearized backbone, were assembled using the Gibson assembly method. All primers for PCR were designed with Gibson-compatible overhangs and ordered from Oligomer. Amplifications were performed with Q5 High-Fidelity DNA Polymerase (New England Biolabs Inc., Boston, USA), and annealing temperatures were determined using NEB’s Tm Calculator. PCR products were purified from agarose gels using a gel extraction kit (M&N, REF 740609.50). Gibson-assembled plasmids were transformed into chemically competent cells by heat shock, followed by overnight incubation on LB agar containing antibiotics for selection. Positive colonies were inoculated into LB broth with antibiotics for plasmid propagation. Plasmids were purified using the GeneJET Plasmid Miniprep Kit (Thermo Fisher Scientific, K0502). Sequences of primers and gene fragments are provided in Tables S1 and S2, with cloning maps in Figure S2. Final plasmid sequences were confirmed by Next Generation Sequencing (NGS) (Intergen, Figure S3).

### Expression of Native and Bispecific Antibodies in CHO Expression System and Stable Pool Development

After sequence verification, bacterial colonies were cultured in LB with antibiotics, mixed 1:1 with 50% glycerol, and stored at –80°C. For plasmid prep, 50 mL LB cultures were grown and processed using the NucleoBond Xtra Midi Kit (Macherey-Nagel). Monospecific and bispecific antibodies were expressed in CHO cells cultured in EX-CELL CHO Fusion medium with 4% L-glutamine. Cells were expanded in T-75 flasks and later cultured in shaking flasks at 37°C, 5% CO₂, 125–130 rpm. Cells were passaged twice weekly at a density of 0.3 × 10^6 cells/mL. Cryopreservation was performed using 7% DMSO and a gradual freezing method before storage in liquid nitrogen.

#### Stable Cell Line Development

CHO® GS⁻/⁻ cells were thawed from cryostorage and seeded at a final density of 0.5 × 10⁶ cells/mL in EX-CELL® CD CHO Fusion Growth Medium supplemented with 6 mM L-glutamine, used for both transfection and recovery. Each T-25 cm² suspension flask was filled with 5.0 mL of complete medium under aseptic conditions. Electroporation cuvettes (0.4 cm gap) were pre-chilled on ice. A total of 4 × 10⁶ cells in 800 µL and 40 µg of purified plasmid DNA were transferred to a cuvette and electroporated using the Gene Pulser MXcell system (Bio-Rad) with manual settings (300 V, 950 µF, exponential decay). Following electroporation, 600 µL of the cell mixture was added to the T-25 cm² flask and incubated. At 24 hours post-transfection, cells were transferred to a sterile 15 mL conical tube, centrifuged at 220 rcf for 5 minutes at 15–25°C, and resuspended in 10.0 mL EX-CELL® CD CHO Fusion Medium without L-glutamine (GS selection medium). The suspension was transferred to a T-75 cm² flask for stable pool selection.

Full medium exchange was performed weekly. Cultures were incubated at 37°C with 5% CO₂ and passaged biweekly to maintain 0.3 × 10⁶ viable cells/mL. Cell counts were recorded weekly.

### Purification of Native and Bispecific Antibodies

For protein purification, plasmids that had been cloned and verified were transfected into CHO cells by electroporation. After generating a stable pool, purification steps were initiated with an initial work volume of 150 ml.

#### Capto L affinity chromatography

In this purification method, the samples were first diluted with binding buffer, and the column was washed with degassed water to a volume equal to 1 column volume (CV). For equilibration of the column, 50 mM Tris base, 3 M NaCl, pH 6.5, at 1 mL/min was employed for equilibration for at least 5 CV. After that, the samples were loaded. Two elution buffers were used. Elution buffer A (20 mM sodium citrate, pH: 5) and elution buffer B (20 mM sodium citrate, pH: 2.5) were introduced under linear pH gradient conditions.^58^ At the end of the protocol, the elutions with antibodies were collected in tubes containing 100 μl neutralization buffer (1 M Tris HCl, pH 8.0) per ml fraction. Finally, the column was washed with 2 CV of degassed water, followed by 2 CV of 20% ethanol. For buffer exchange, the protein sample was introduced into a filter column (50 kDa Ultra-4 Centrifugal Filter Units, Amicon). Subsequently, the filter column was subjected to centrifugation at 4000 g for 45 minutes at 4°C. This step was repeated until all of the sample was passed into the filter column. Then, PBS was added to the filter column, and the process was repeated at 4°C until the buffer exchange was completed. At last, the purified IgG samples were filtered using a 0.2 µm filter.

#### SEC Analysis

In this study, bispecific antibodies and parental monoclonal antibodies were subjected to size exclusion chromatography (SEC). After transfection, the antibodies in the cell culture supernatant were collected and filtered through a 0.2 µm PES membrane (Thermo Scientific). The protein purification was achieved using a HiPrep 16/60 Sephacryl S-200 HR column (Cytiva) with FPLC (ÄKTA start protein purification system), following the manufacturer’s prescribed protocol. Degassed and pre-filtered, all solutions were prepared. For equilibration of the column, degassed and filtered water was passed through at a flow rate of 0.5 mL/min for one-half column volume. 0.05 M sodium phosphate, 0.15 M NaCl, pH 7.2, at 1 mL/min were employed for equilibration. As part of the regular cleaning process, the column was washed with 0.2 M NaOH at a flow rate of 0.5 mL/min to eliminate most proteins that are non-specifically bound to the medium. After this cleaning step, it is essential to promptly re-establish the column by equilibrating it with a minimum of two column volumes of buffer. Hence, this wash step was continued until the UV baseline had stabilized. At the end of the protocol, the elutions having antibodies were collected in tubes (Figure S4). For buffer exchange, the same protocol was applied. Finally, the column was washed with 4 CV degassed water, followed by 4 CV 20% ethanol, and then was stored at room temperature. Based on the information in the protocol, it was noted that the elutions of the first peak collected IgG. A western blot was performed to determine the three peaks, and it was observed that only the first peak contained IgG.

Detailed Protein A and MabSelect purification protocols, beyond the two methods described above, are provided in the Supplementary Material and Methods section.

### SDS-PAGE and Western Blotting

For western blot analysis, protein samples were heat-treated at 95 °C with 1× SDS loading dye. Gel electrophoresis was conducted using an 8% SDS polyacrylamide gel. Following loading of the samples into the gel and completion of the gel run, the gel’s contents were transferred to a PVDF membrane using the Trans-Blot Turbo (Bio-Rad) system. For blocking, the PVDF membrane was incubated in 5% skim milk in TBS-T for 1 hour at room temperature. Following blocking, the membrane was incubated in 5% milk in TBS-T with a 1:5000 dilution of horseradish peroxidase (HRP)-conjugated antibodies for 1.5 h at room temperature. When the incubation was complete, the membrane was washed three times with 1x TBS-T for 10 minutes each. Visualization of the membrane was achieved through incubation with ECL substrates (Bio-Rad 170-5060). Images were captured using the Vilber Fusion Solo S system. For the detection of anti-EGFR, anti-HER2, and anti-EGFR/HER2, the secondary antibodies used were conjugated Goat pAb Anti-Human IgG (HRP) (ab98624, Abcam).

### Determination of The Binding Activities of Antibodies Using ELISA

Commercial ELISA Kits were used to determine the binding capabilities of the monoclonal antibodies and bispecific antibodies with EGFR and HER2. The monospecific and bispecific antibody concentrations were quantified using a Trastuzumab mAb-Based ELISA Kit (ImmunoGuide Trastuzumab ELISA, IG-AB105) and a Cetuximab mAb-Based ELISA Kit (ImmunoGuide Cetuximab ELISA, IG-AB112), following the recommended protocols provided by the manufacturers.

### Sandwich ELISA

A dual-binding ELISA was established to assess the binding activity of the anti-EGFR/HER2 bispecific antibody. In this method, 96-well plates were coated with 1 ng/µl Anti-Cetuximab mouse monoclonal antibody, which was prepared with PBST, incubated overnight at 4 °C, and subsequently blocked with blocking solution, comprising 5% bovine serum albumin (BSA) in PBS plus 0.1% Tween-20 (PBST) for 1 hour at 37 °C. After three washes with PBST, the plate was exposed to a 100 µl sample volume of bispecific antibodies and negative controls for 1 hour at 37 °C, and then washed again with PBST. After the washing step, 100 µl anti-Trastuzumab (HRP) was introduced separately. Following an hour incubation, the determination of binding affinity was achieved by measuring absorbance at 450 nm using a microplate reader (SpectraMax M5), after incubation with the 3,3′,5,5′-tetramethylbenzidine (TMB) substrate reagent, and the reaction was stopped by adding Stop solution (1 N Hydrochloric acid (HCl)).

### Circular Dichroism (CD)

Circular dichroism (CD, JASCO J-815, Tokyo, Japan) studies were conducted to observe alterations in secondary structures between bispecific antibodies and native monospecific antibodies. Each protein was prepared at a concentration of 200 µg/mL in 2 mM potassium phosphate buffer, pH 7.4, for the experiments. The three channels selected for analysis included CD, voltage, and absorbance. The wavelength range was set from 200 to 240 nm, with a 4-second digital integration time (D.I.T.), a 1 nm bandwidth, standard sensitivity, 100 nm/min scanning speed, and 3-repeat accumulation modes.

### Immunocytochemistry (ICC)

15 mm coverslips were treated with ethanol overnight. Then, the coverslips were dried under the hood and placed into the wells of 24-well plates. Placed coverslips were treated with collagen (C8919, Sigma-Aldrich) for 15 minutes and then dried after the collagen was removed. Caco-2, JIMT-1, and CHOK-1 cells were seeded at densities of 2 × 10^5^ cells, 2 × 10^5^ cells, and 1 × 10^5^ cells per well, respectively, onto coverslips and incubated at 37 °C for 24 hours. Then, cells were fixed with ice-cold 10% formalin and washed three times with ice-cold PBS. For permeabilization, only cells to be treated with the β-Actin antibody were incubated in 0.2% Triton X-100 for 10 minutes at room temperature, while gently shaking. Then, cells were blocked with 2% BSA in PBS for 1 hour while gently shaking at room temperature. After blocking, cells were incubated overnight at 4 °C with primary antibodies diluted with 2% BSA in PBS (anti-HER2 (1:500), anti-EGFR (1:500), anti-EGFR/HER2 (1:500), and β-Actin (8H10D10) Mouse mAb (1:250, 3700, Cell Signaling)) and washed with 0.1% PBST three times for 15 minutes while gently shaking. Then, cells were incubated with secondary antibodies diluted with 2% BSA in PBS (Goat anti-Mouse IgG (H+L) Secondary Antibody, FITC (1:100, 31569, Thermo Fisher Scientific) and Alexa Fluor® 488 anti-human IgG Fc Antibody (1:1000, 410705, Biolegend)) for 2 hours at room temperature and washed with 0.1% PBST three times for 15 minutes while gently shaking. Removed coverslips were mounted with Ibidi Mounting Medium With DAPI (50011, Ibidi). Slides were imaged with a Leica Confocal Microscope.

### *In Vitro* Cytotoxicity Tests

Caco-2 cells were maintained in Minimum Essential Medium Eagle (MEM) supplemented with 15% fetal bovine serum (FBS), 1% Penicillin/Streptomycin (Pen/Strep), and 1% L-glutamine. After thawing, the contents of the stock vial were diluted in 13 mL of fresh medium and then centrifuged at 2500 rpm for 5 minutes. The supernatant was subsequently discarded. The cell pellet was resuspended in 1 mL of fresh medium, transferred to a T25 flask containing 2 mL of media, and incubated at 37°C with 5% CO₂. At 80–90% confluency, the cells were trypsinized with 2 mL of trypsin, incubated for 5 minutes, and then neutralized with 4 mL of medium. Following centrifugation, cells were resuspended in fresh medium and transferred to new flasks. Cell counts were performed routinely, and cells were screened for mycoplasma contamination. JIMT-1 cells were cultured in low-glucose DMEM containing 15% FBS, 1% Pen/Strep, and 1% L-glutamine, with similar subculturing procedures.

#### Cell Proliferation Assay MTT

Caco-2, JIMT-1, and CHOK-1 cells were seeded into 96-well plates at a density of 9 × 10³ cells (about 30% confluency) per well and incubated at 37 °C for 24 hours. Epidermal growth factor (EGF) was added to a final concentration of 10 nmol/L, and Caco-2 cells were further incubated for 24 hours. The EGF treatment was exclusively applied to Caco-2 cells. Subsequently, the specified antibodies were introduced for 24 hours. After this, the cells were treated with MTT solution (5 mg/mL in PBS) for 4 hours at 37 °C to allow the formation of blue formazan crystals in metabolically active cells. The crystals were then solubilized using DMSO, and cell proliferation was quantified by measuring the absorbance at 570 nm using a microplate reader (Biotek Synergy HT Microplate Reader). The findings were presented as a percentage of MTT reduction compared to the absorbance of the control cells.

#### 3D Cell Culture

A 1.5% high-purity agarose solution was prepared in PBS and sterilized by autoclaving for use in mammalian cell spheroid culture. A total of 100 µL of sterilized agarose was dispensed into each well of a 96-well plate and allowed to solidify for 2 hours at room temperature. Trypsinized cells were resuspended in fresh culture medium following centrifugation and seeded onto the solidified agarose at a density of 7,500 cells/well. Cultures were incubated until spheroids were formed. The spheroid medium did not contain FBS for any cell line. For Caco-2 cells, spheroid formation was additionally evaluated in the presence or absence of EGF supplementation. Spheroid formation required approximately 5 days for Caco-2, 3 days for JIMT-1, and 2 days for CHO-K1 cells. Following spheroid formation, antibodies were added to the wells at final concentrations of 1 μg/mL and 10 μg/mL. Cell viability and cytotoxicity were monitored using the CellTiter-Glo® 3D Cell Viability Assay (Promega) according to the manufacturer’s instructions. Also, Zeiss fluorescence microscope was utilized.

#### Quantitative RT-PCR analysis

To assess apoptosis via the mRNA expression levels of Bax and Bcl-2 using quantitative RT-PCR, 30-40% confluent cancer cell cultures (Caco-2 and JIMT-1) were cultured in 48-well plates for 24 hours of incubation. After a 24-hour incubation, monospecific and bispecific antibodies were added at a concentration of 10 µg/mL for each, and this treatment was continued for an additional 18 hours. For Caco-2 cell culture was applied two different conditions as with and without EGF induction. As a control, a corresponding amount of culture medium was used in place of the compound. RNA was extracted using the Monarch® Total RNA Miniprep Kit (NEB, T2010S). RNA purity was verified by 260/280 nm absorbance, and cDNA synthesis was performed using LunaScript® RT SuperMix Kit (E3010). Template cDNA (40 ng per sample) was used in qPCR with Luna® Universal qPCR Master Mix on a Biorad thermocycler. To ensure accuracy, all experiments were conducted in triplicate. Reactions included an initial denaturation (94°C, 3 min), 40 cycles of 95°C (60 s), 60°C (30 s), and a melt curve step. Sequences corresponding to the qRT-PCR primers are listed in Supplementary Table S3. Gene expression levels for Bax, Bcl-2, and Actin were analyzed using the 2^-ΔΔCT method,^59^ allowing for a relative quantification of mRNA expression across different conditions. Using this method, it was determined whether both cell lines expressed EGFR and HER2 genes (Figure S5).

#### Antibody-dependent Cell-mediated Cytotoxicity Assay

Two target cell lines, Caco-2 and JIMT-1, along with the human monocytic cell line THP-1, were cultured following the supplier’s recommendations. All required media compositions and conditions were previously detailed. All cells were maintained in a 5% CO₂-enriched atmosphere at 37 °C. Caco-2 and JIMT-1 cells were seeded into 96-well plates at 30% confluency in 2% FBS medium and labeled with 1 µM calcein-AM (Thermo Fisher) at 37°C for 30 minutes in the dark. Excess dye was removed by washing with PBS, and labeling efficiency was confirmed via fluorescence microscopy. CD14⁺ THP-1 effector cells were washed and resuspended in RPMI-1640 with 2% FBS, then added at an effector-to-target ratio of 5:1 (total 100 µL/well). Cells were treated with 10 µg/mL of bispecific or monospecific antibodies and incubated for 4 hours at 37°C. Control wells included spontaneous release (target cells alone) and maximum release (1% Triton X-100). Post-incubation, 100 µL of supernatant was transferred for fluorescence analysis. Fluorescence intensity was measured using a microplate reader with excitation and emission wavelengths of 485 nm and 520 nm, respectively. The percentage of calcein release was calculated using the following formula: *Calcein Release (%) = (Experimental Release−Spontaneous Release) / Maximum Release−Spontaneous Release)×100*

### Statistical Analysis

The figures present data drawn using GraphPad Prism 8. Statistical analysis was carried out using the software tools. Significance levels indicated in the figures were determined through calculations using unpaired t-tests and one-way ANOVA. Each condition was performed in triplicate, and data were presented as the mean ± standard deviation. Statistical significance is represented by *, **, ***, and **** corresponding to p < 0.05, 0.01, 0.001, and 0.0001, respectively. Changes in the trial track were recorded. Visual representations were produced with Biorender.com.

## Supporting Information

Gene sequences and primers used; Gel images of Bax and Bcl-2 PCR products; FPLC-based antibody purification workflow; EGFR/HER2 expression in cell lines; Plasmid maps and corresponding sequencing results generated in this study.

## Author Information

### Authors

**Senem Sen -** *UNAM─Institute of Materials Science and Nanotechnology, National Nanotechnology Research Center, Bilkent University, Ankara 06800, Turkey*

**Aslı Semerci -** *UNAM─Institute of Materials Science and Nanotechnology, National Nanotechnology Research Center, Bilkent University, Ankara 06800, Turkey*

**Melis Karaca -** *UNAM─Institute of Materials Science and Nanotechnology, National Nanotechnology Research Center, Bilkent University, Ankara 06800, Turkey*

**Recep Erdem Ahan -** *UNAM─Institute of Materials Science and Nanotechnology, National Nanotechnology Research Center, Bilkent University, Ankara 06800, Turkey*

**Cemile Elif Ozcelik -** *UNAM─Institute of Materials Science and Nanotechnology, National Nanotechnology Research Center, Bilkent University, Ankara 06800, Turkey*

**Elif Duman -** *Synbiotik Biotechnology and Biomedical Technology, Cankaya, Ankara, Turkey* **Behide Saltepe -** *UNAM─Institute of Materials Science and Nanotechnology, National Nanotechnology Research Center, Bilkent University, Ankara 06800, Turkey*

**Ebru Sahin Kehribar -** *Synbiotik Biotechnology and Biomedical Technology, Cankaya, Ankara, Turkey*

**Ebru Aras -** *UNAM─Institute of Materials Science and Nanotechnology, National Nanotechnology Research Center, Bilkent University, Ankara 06800, Turkey*

**Eray Ulas Bozkurt -** *UNAM─Institute of Materials Science and Nanotechnology, National Nanotechnology Research Center, Bilkent University, Ankara 06800, Turkey*

**Urartu Ozgur Safak Seker -** *UNAM─Institute of Materials Science and Nanotechnology, National Nanotechnology Research Center, Bilkent University, Ankara 06800, Turkey*

### Author Contributions

US designed the study, analyzed the results, secured funding; SS; run the protein expression studies, molecular clonings of the final designs, and did the molecular characterizations and cell assays; EU gave feedback on the design and results, secured funding; ED, ESK, RA, BS constructed the initial genetic designs, EA, EUB run protein expression studies, AS and MK did protein expression, AS also did molecular characterizations, spheroid experiments, CEO and MK did ICC assay.

## Acknowledgement

This research received partial funding from the Scientific and Technological Research Council of Turkey (TUBITAK), Grant Numbers: 119C054 and 217S118.

