## Supplemental data for "Developing Anti-EGFR/Anti-HER2 Bifunctional Antibody for Solid Tumors by Protein Engineering"

**Contents**

*Supplementary Table S1*

*Supplementary Table S2*

*Supplementary Table S3*

*Supplementary Figure S1*

*Supplementary Figure S2*

*Supplementary Figure S3*

*Supplementary Figure S4*

*Supplementary Figure S5*

*Supplementary Figure S6*

*Supplementary Figure S7*

*Supplementary Figure S8*

*Supplementary Material and Methods*

*Supplementary References*

**Table S1.** Gene sequences used in the study (from 5' to 3')

| Sequence Name | Sequence (5' to 3') |
|---|---|
| CMV enhancer | GACATTGATTATTGACTAGTTATTAATAGTAATCAATTACGGG GTCATTAGTTCATAGCCCATATATGGAGTTCCGCGTTACATA ACTTACGGTAAATGGCCCGCCTGGCTGACCGCCCAACGACC CCCGCCCATTGACGTCAATAATGACGTATGTTCCCATAGTAA CGCCAATAGGGACTTTCCATTGACGTCAATGGGTGGAGTATT TACGGTAAACTGCCCACTTGGCAGTACATCAAGTGTATCATA TGCCAAGTACGCCCCCTATTGACGTCAATGACGGTAAATGG CCCGCCTGGCATTATGCCCAGTACATGACCTTATGGACTTT CCTACTTGGCAGTACATCTACGTATTAGTCATCGCTATTACCA TG |
| CMV promoter | GTGATGCGGTTTTGGCAGTACATCAATGGGCGTGGATAGCG GTTTGACTCACGGGGATTTCCAAGTCTCCACCCCATTGACGT CAATGGGAGTTTGTTTTGGCACCAAAATCAACGGGACTTTCC AAAATGTCGTAACAACTCCGCCCCATTGACGCAAATGGGCG GTAGGCGTGTACGGTGGGAGGTCTATATAAGCAGAGCT |
| Kozak sequence | GCCGCCACCATGG |
| Light chain signal sequence | ATGGGCTGGAGCTGTATCATTCTGTTTCTGGTGGCCACAGC CACTGGCGTGCACAGC |
| Anti-EGFR native light chain | GCCACCATGGGCTGGAGCTGTATCATTCTGTTTCTGGTGGC CACAGCCACTGGCGTGCACAGCGACATACTCCTCACCCAGT CTCCAGTAATTCTGAGCGTTAGTCCCGGCGAGCGTGTGAGT TTTTCCTGCCGCGCATCACAGTCCATAGGAACAAATATACAT TGGTATCAGCAGAGAACCAACGGCTCCCCTCGCCTGCTCAT AAAGTACGCTTCAGAGTCCATATCAGGGATACCATCCCGGTT TTCAGGATCTGGCAGTGGTACTGACTTTACCTTGTCAATTAAT TCAGTTGAGAGCGAAGACATAGCAGATTATTATTGCCAGCAA AACAATAACTGGCCCACCACCTTTGGAGCTGGAACAAAGCT GGAGTTGAAGCGAACAGTCGCAGCCCCTTCAGTCTTCATCTT TCCCCCCTCAGACGAGCAACTTAAATCAGGCACTGCCTCCG TAGTGTGTCTTCTTAACAACTTCTATCCACGGGAAGCTAAAG TCCAGTGGAAGGTGGATAATGCTTTGCAATCTGGGAATAGTC AAGAATCAGTAACAGAGCAGGACAGTAAAGACTCAACCTACA GCTTGAGTTCAACACTTACACTCTCTAAGGCTGATTATGAGA AGCACAAAGTCTATGCCTGTGAGGTAACCCACCAGGGTCTG TCATCACCAGTGACAAAGAGCTTCAACAGAGGCGAGTGTTG ATGA |
| Anti-HER2 native light chain | ATGGGCTGGAGCTGTATCATTCTGTTTCTGGTGGCCACAGC CACTGGCGTGCACAGCGACATACAAATGACTCAGTCCCCAT CAAGTCTCAGTGCCAGTGTAGGTGACCGAGTAACTATTACTT GTAGGGCCTCCCAAGACGTAAACACAGCAGTGGCATGGTAT CAACAGAAGCCTGGAAAGGCACCCAAATTGCTTATATACTCT GCCAGTTTTCTTTACAGCGGCGTCCCCAGCCGATTCTCAGGT TCTAGGAGCGGAACTGACTTCACTCTCACAATTTCTTCTTTG CAACCTGAGGATTTTGCTACTTATTACTGCCAACAGCATTACA CCACTCCACCCACCTTTGGTCAGGGCACTAAGGTCGAAATTA AGCGGACTGTCGCTGCTCCTTCAGTCTTCATTTTTCCCCCTT CTGACGAACAACTCAAATCAGGCACTGCCAGTGTCGTATGTT TGTTGAATAACTTTTATCCCCGAGAAGCAAAAGTGCAATGGA AGGTAGACAATGCACTGCAAAGCGGTAACAGCCAGGAAAGT |

| | |
|---|---|
| | GTGACAGAACAGGATTCAAAGGATAGCACATATTCCCTCTCC TCCACTCTGACCCTTTCTAAAGCCGACTACGAGAAACACAAA GTTTACGCTTGTGAGGTTACACACCAGGGATTGTCTAGCCCT GTGACAAAAAGCTTCAACAGAGGCGAGTGTTGATGA |
| Anti- HER2 Crossmab light chain | GAGGTTCAGTTGGTCGAGTCAGGTGGTGGTCTTGTCCAACC AGGCGGTTCATTGCGTTTGAGTTGTGCCGCTTCCGGTTTTAA CATAAAGGATACTTATATACATTGGGTCAGACAGGCACCAGG TAAGGGGCTGGAGTGGGTTGCAAGGATATATCCTACAAACG GTTACACAAGGTATGCTGACTCTGTTAAGGGCCGCTTTACCA TAAGCGCTGACACCTCAAAAAATACTGCTTATTTGCAAATGAA CAGCCTTAGAGCTGAAGATACCGCAGTATATTATTGCTCAAG GTGGGGTGGTGACGGATTCTATGCCATGGACTACTGGGGCC AAGGTACACTTGTTACTGTTTCTTCC |
| Heavy chain signal sequence | GCCACCATGGAGCTCGGCCTCAGCTGGGTCTTTCTGGTGGT CATTCTGGAGGGCGTGCAATGT |
| Anti-EGFR native heavy chain | ATGGAGCTCGGCCTCAGCTGGGTCTTTCTGGTGGTCATTCT GGAGGGCGTGCAATGTCAAGTGCAGCTGAAGCAGAGCGGA CCCGGCCTCGTGCAACCTTCCCAGTCTCTGAGCATCACATG TACTGTCAGCGGCTTCTCTCTGACTAACTACGGCGTGCATTG GGTGAGGCAGTCCCCCGGCAAGGGACTGGAGTGGCTGGGC GTCATCTGGTCCGGCGGCAACACTGACTACAACACACCTTTC ACATCTAGGCTGAGCATTAATAAGGACAATTCCAAATCCCAA GTGTTTTTCAAGATGAACTCTCTGCAGTCCAATGACACTGCT ATCTACTACTGCGCCAGAGCCCTCACATACTACGACTACGAG TTCGCTTACTGGGGCCAAGGCACTCTGGTGACTGTGTCCGC CGCCAGCACTAAAGGCCCAAGCGTGTTCCCTCTGGCTCCTA GCTCCAAAAGCACTTCCGGCGGAACTGCTGCTCTGGGCTGT CTCGTGAAGGACTACTTCCCAGAACCAGTGACAGTGAGCTG GAATAGCGGCGCTCTGACTAGCGGAGTCCATACATTTCCAG CCGTGCTCCAGAGCAGCGGCCTCTACAGCCTCAGCTCCGTG GTGACAGTGCCAAGCAGCTCCCTCGGCACACAGACATACAT CTGCAATGTCAACCACAAGCCTTCCAACACTAAGGTGGACAA GAGGGTCGAGCCTAAGAGCTGTGACAAGACACACACTTGTC CTCCATGCCCAGCTCCAGAGCTGCTCGGCGGACCATCCGTC TTTCTGTTTCCTCCTAAGCCAAAGGACACACTGATGATCTCTA GGACTCCAGAGGTGACATGCGTCGTGGTGGACGTCAGCCAC GAAGACCCAGAGGTCAAGTTCAACTGGTACGTCGACGGAGT CGAAGTCCACAACGCCAAGACTAAGCCTAGGGAGGAACAGT ACAACTCCACTTATAGGGTCGTGTCCGTCCTCACTGTGCTGC ACCAAGATTGGCTCAACGGCAAGGAGTACAAGTGTAAGGTG TCCAATAAGGCTCTGCCAGCCCCTATCGAGAAGACAATCAG CAAGGCTAAGGGCCAGCCAAGGGAGCCACAAGTCTATACAC TCCCACCATCTAGGGAAGAGATGACAAAGAACCAAGTGTCTC TGACTTGTCTGGTGAAGGGATTCTACCCAAGCGATATTGCCG TGGAATGGGAGAGCAACGGACAGCCAGAGAACAACTACAAG ACTACACCACCAGTGCTGGACTCCGATGGCAGCTTCTTCCTC TACAGCAAGCTCACTGTCGACAAGTCTAGGTGGCAGCAAGG CAACGTGTTCAGCTGTTCCGTCATGCATGAAGCTCTGCACAA CCACTACACACAGAAGAGCCTCAGCCTCAGCCCCGGCAAGT GA |

| | |
|---|---|
| Anti-HER2 native heavy chain | ATGGAGCTCGGCCTCAGCTGGGTCTTTCTGGTGGTCATTCT<br>GGAGGGCGTGCAATGTGAGGTTCAGTTGGTCGAGTCAGGTG<br>GTGGTCTTGTCCAACCAGGCGGTTCATTGCGTTTGAGTTGTG<br>CCGCTTCCGGTTTTAACATAAAGGATACTTATATACATTGGGT<br>CAGACAGGCACCAGGTAAGGGGCTGGAGTGGGTTGCAAGG<br>ATATATCCTACAAACGGTTACACAAGGTATGCTGACTCTGTTA<br>AGGGCCGCTTTACCATAAGCGCTGACACCTCAAAAAATACTG<br>CTTATTTGCAAATGAACAGCCTTAGAGCTGAAGATACCGCAG<br>TATATTATTGCTCAAGGTGGGGTGGTGACGGATTCTATGCCA<br>TGGACTACTGGGGCCAAGGTACACTTGTTACTGTTTCTTCCG<br>CATCCACCAAAGGCCCATCAGTATTTCCACTCGCTCCCAGCA<br>GTAAGTCTACATCAGGAGGAACAGCTGCCCTTGGCTGTCTC<br>GTCAAAGACTACTTTCCTGAACCCGTAACCGTTAGTTGGAAC<br>TCCGGGGCATTGACAAGCGGTGTCCATACTTTCCCCGCAGT<br>CTTGCAGTCCAGTGGCTTGTATTCTCTCTCTAGTGTCGTAAC<br>AGTCCCATCATCTTCACTTGGCACCCAGACCTATATCTGCAA<br>TGTAAATCACAAACCATCTAATACTAAGGTAGATAAAAAAGTC<br>GAGCCTAAGAGCTGCGACAAAACTCACACTTGCCCCCCTTG<br>TCCAGCACCTGAGCTGTTGGGCGGCCCAAGCGTCTTTCTTTT<br>CCCACCAAAGCCAAAGGACACCTTGATGATTTCTAGGACACC<br>AGAAGTGACTTGTGTAGTCGTAGACGTTTCACACGAAGATCC<br>TGAAGTAAAATTTAATTGGTACGTCGATGGGGTAGAGGTGCA<br>CAATGCCAAAACTAAGCCCAGAGAAGAACAATACAACAGCAC<br>ATACAGAGTTGTCAGTGTATTGACAGTTCTCCATCAAGATTG<br>GCTCAACGGTAAAGAGTATAAATGTAAGGTCAGTAACAAGGC<br>CCTCCCCGCTCCTATAGAAAAGACAATCTCCAAGGCCAAAG<br>GCCAGCCTAGAGAGCCTCAAGTCTACACACTCCCCCCAAGT<br>AGGGACGAGCTTACAAAAAATCAGGTTTCTCTGACATGCCTG<br>GTAAAGGGGTTCTATCCCAGCGACATTGCCGTCGAATGGGA<br>ATCAAATGGTCAGCCCGAAAATAACTATAAAACCACACCACC<br>CGTTCTCGATTCCGACGGAAGCTTCTTTCTCTACTCAAAGCT<br>TACAGTGGACAAGAGTCGTTGGCAGCAGGGCAATGTTTTTA<br>GTTGCTCTGTGATGCACGAAGCACTGCACAACCATTATACCC<br>AAAAAAGTCTTAGCCTCAGCCCCGGCAAGTGATGA |
| Anti- HER2 hole-mutated heavy chain | GCCGCCACCATGGAGCTCGGCCTCAGCTGGGTCTTTCTGGT<br>GGTCATTCTGGAGGGCGTGCAATGTGACATACAAATGACTCA<br>GTCCCCATCAAGTCTCAGTGCCAGTGTAGGTGACCGAGTAA<br>CTATTACTTGTAGGGCCTCCCAAGACGTAAACACAGCAGTGG<br>CATGGTATCAACAGAAGCCTGGAAAGGCACCCAAATTGCTTA<br>TATACTCTGCCAGTTTTCTTTACAGCGGCGTCCCCAGCCGAT<br>TCTCAGGTTCTAGGAGCGGAACTGACTTCACTCTCACAATTT<br>CTTCTTTGCAACCTGAGGATTTTGCTACTTATTACTGCCAACA<br>GCATTACACCACTCCACCCACCTTTGGTCAGGGCACTAAGGT<br>CGAAATTAAGTCTTCCGCATCCACCAAAGGCCCATCAGTATT<br>TCCACTCGCTCCCAGCAGTAAGTCTACATCAGGAGGAACAG<br>CTGCCCTTGGCTGTCTCGTCAAAGACTACTTTCCTGAACCCG<br>TAACCGTTAGTTGGAACTCCGGGGCATTGACAAGCGGTGTC<br>CATACTTTCCCCGCAGTCTTGCAGTCCAGTGGCTTGTATTCT<br>CTCTCTAGTGTCGTAACAGTCCCATCATCTTCACTTGGCACC<br>CAGACCTATATCTGCAATGTAAATCACAAACCATCTAATACTA<br>AGGTAGATAAAAAAGTCGAGCCTAAGAGCTGCGACAAAACTC |

| | |
|---|---|
| | ACACTTGCCCCCCTTGTCCAGCACCTGAGCTGTTGGGCGGC CCAAGCGTCTTTCTTTTCCCACCAAAGCCAAAGGACACCTTG ATGATTTCTAGGACACCAGAAGTGACTTGTGTAGTCGTAGAC GTTTCACACGAAGATCCTGAAGTAAAATTTAATTGGTACGTC GATGGGGTAGAGGTGCACAATGCCAAAACTAAGCCCAGAGA AGAACAATACAACAGCACATACAGAGTTGTCAGTGTATTGAC AGTTCTCCATCAAGATTGGCTCAACGGTAAAGAGTATAAATG TAAGGTCAGTAACAAGGCCCTCCCCGCTCCTATAGAAAAGAC AATCTCCAAGGCCAAAGGCCAGCCTAGAGAGCCTCAAGTCT GCACACTCCCCCCAAGTAGGGACGAGCTTACAAAAAATCAG GTTTCTCTGTCCTGCGCCGTAAAGGGGTTCTATCCCAGCGA CATTGCCGTCGAATGGGAATCAAATGGTCAGCCCGAAAATAA CTATAAAACCACACCACCCGTTCTCGATTCCGACGGAAGCTT CTTTCTCGTTTCAAAGCTTACAGTGGACAAGAGTCGTTGGCA GCAGGGCAATGTTTTTAGTTGCTCTGTGATGCACGAAGCACT GCACAACCATTATACCCAAAAAAGTCTTAGCCTCAGCCCCGG CAAGTGATGA |
| Anti-EGFR knob-mutated heavy chain | ATGGAGCTCGGCCTCAGCTGGGTCTTTCTGGTGGTCATTCT GGAGGGCGTGCAATGTCAAGTGCAGTTGAAGCAATCAGGTC CCGGCTTGGTGCAGCCAAGTCAAAGTCTGTCTATTACTTGCA CAGTGAGCGGCTTTTCTCTCACCAATTACGGCGTCCACTGG GTTCGGCAAAGTCCAGGAAAGGGACTTGAATGGCTGGGGGT CATCTGGAGTGGGGGTAATACAGATTACAATACCCCATTCAC CTCTAGACTGAGTATAAATAAGGATAATTCCAAATCCCAAGT GTTCTTTAAGATGAACTCACTTCAGAGTAATGATACAGCTATA TACTATTGTGCCCGAGCCCTGACATACTATGACTACGAGTTT GCCTACTGGGGACAAGGAACACTGGTCACCGTCAGCGCCG CAAGCACCAAAGGCCCCAGTGTATTTCCACTTGCCCCAAGTA GTAAAAGTACTTCAGGTGGAACAGCCGCTTTGGGCTGCCTG GTAAAGGATTACTTCCCCGAACCAGTTACTGTGTCTTGGAAT AGCGGTGCTCTTACTTCCGGCGTCCACACTTTCCCCGCTGT GCTCCAATCTTCAGGCTTGTATTCCCTGAGCAGTGTTGTAAC TGTCCCATCAAGTTCATTGGGAACCCAAACTTACATATGTAAT GTGAATCATAAGCCTAGCAACACTAAAGTGGATAAGAAAGTG GAACCAAAGTCTTGCGACAAGACTCACACTTGCCCCCCTTGT CCAGCACCTGAGCTGTTGGGCGGCCCAAGCGTCTTTCTTTT CCCACCAAAGCCAAAGGACACCTTGATGATTTCTAGGACACC AGAAGTGACTTGTGTAGTCGTAGACGTTTCACACGAAGATCC TGAAGTAAAATTTAATTGGTACGTCGATGGGGTAGAGGTGCA CAATGCCAAAACTAAGCCCAGAGAAGAACAATACAACAGCAC ATACAGAGTTGTCAGTGTATTGACAGTTCTCCATCAAGATTG GCTCAACGGTAAAGAGTATAAATGTAAGGTCAGTAACAAGGC CCTCCCCGCTCCTATAGAAAAGACAATCTCCAAGGCCAAAG GCCAGCCTAGAGAGCCTCAAGTCTACACACTCCCCCCATGC AGGGACGAGCTTACAAAAAATCAGGTTTCTCTGTGGTGCTTG GTAAAGGGGTTCTATCCCAGCGACATTGCCGTCGAATGGGA ATCAAATGGTCAGCCCGAAAATAACTATAAAACCACACCACC CGTTCTCGATTCCGACGGAAGCTTCTTTCTCTACTCAAAGCT TACAGTGGACAAGAGTCGTTGGCAGCAGGGCAATGTTTTTA GTTGCTCTGTGATGCACGAAGCACTGCACAACCATTATACCC AAAAAAGTCTT |

**Table S2.** Primers used to amplify the specified region for corresponding cloning (from 5' to 3')

| Primer Name | Sequence (5' to 3') |
|---|---|
| BB-PCR1-F | AGCCTCAGCCCCGGCAAG |
| BB-PCR1-R | GCTGTGCACGCCAGTGGC |
| BB-PCR2-F | AGCTTCAACAGAGGCGAGTGT |
| BB-PCR2-R | ACATTGCACGCCCTCCAGAATG |
| NLF | GCCACAGCCACTGGCGTGCACAGCGACATACAAATGACTCAGTCCCC |
| NLR | TCAACACTCGCCTCTGTTGAAGCTTTTTGTCACAGGGCTAGAC |
| NHF | GTCATTCTGGAGGGCGTGCAATGTGAGGTTCAGTTGGTCGAGT |
| NHR | TTTGTCGCAGCTCTTAGGCTC |
| Fcc-F | GAGCCTAAGAGCTGCGACAAAACTCACACTTGCCCCC |
| Fcc-R | TCATCACTTGCCGGGGCTGAGGCTAAGACTTTTTTGGGTATAATGGT |
| SP-F | ATGGGCTGGAGCTGTATC |
| SP-R | CCGGTATTGTCTCCTTCCGTG |
| SP-PCR2-F | GTCTTAGCCTCAGCCCCGGCAAGTGATGATTAATTAACTCGAGTACGATAGCCG |
| SP-PCR2-R | CACCAGAAACAGAATGATACAGCTCCAGCCCATGGTGGCGGCTTC |

**Table S3.** Primers used to amplify the specified region for corresponding qRT-PCR amplification of target genes. (from 5' to 3')

| Target Gene | Primer Sequences |
|---|---|
| Bax F<br>Bax R | 5'- CCCGAGAGGTCTTTTTCCGAG -3'<br>5'- CCAGCCCATGATGGTTCTGAT -3' |
| Bcl-2 F<br>Bcl-2 R | 5'- TGGAAAGCGTAGACAAGGAGA -3'<br>5'- TGCTGCATTGTTCCCGTAGA -3' |
| EGFR F<br>EGFR R | 5'- AGGCACGAGTAACAAGCTCAC -3'<br>5'- ATGAGGACATAACCAGCCACC -3' |
| HER2 F<br>HER2 R | 5'- TGCAGGGAAACCTGGAACTC -3'<br>5'- ACAGGGGTGGTATTGTTCAGC -3' |
| Actin F<br>Actin R | 5'- TGACGTGGACATCCGCAAAG -3'<br>5'- CTGGAAGGTGGACAGCGAGG -3' |

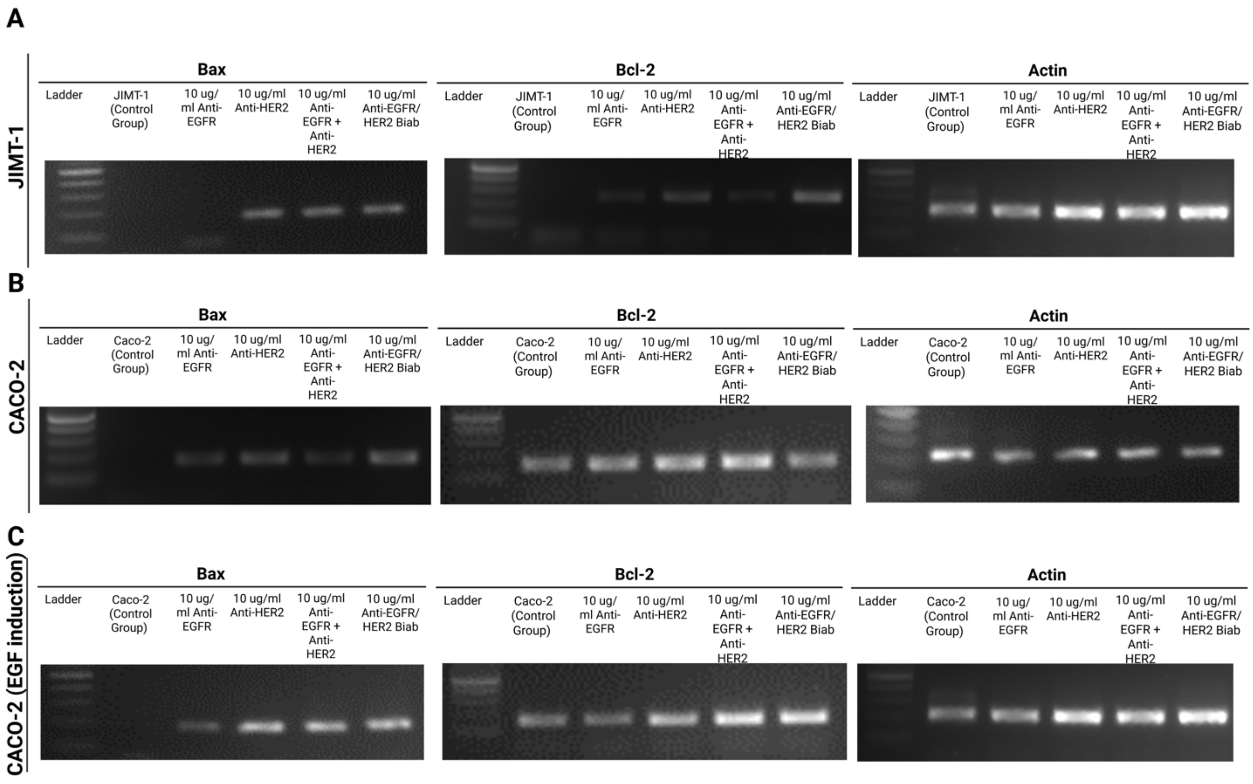


**Figure S1.** (A-C) Cropped gel images showing PCR products of Bax and Bcl-2 gene amplifications. All samples originate from a single experiment, and gels were processed simultaneously.

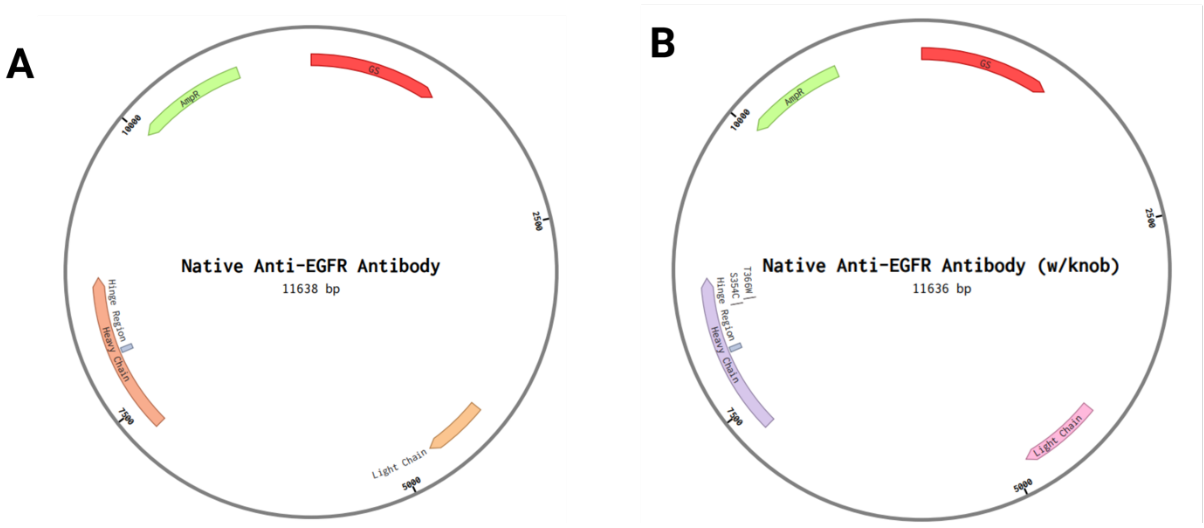

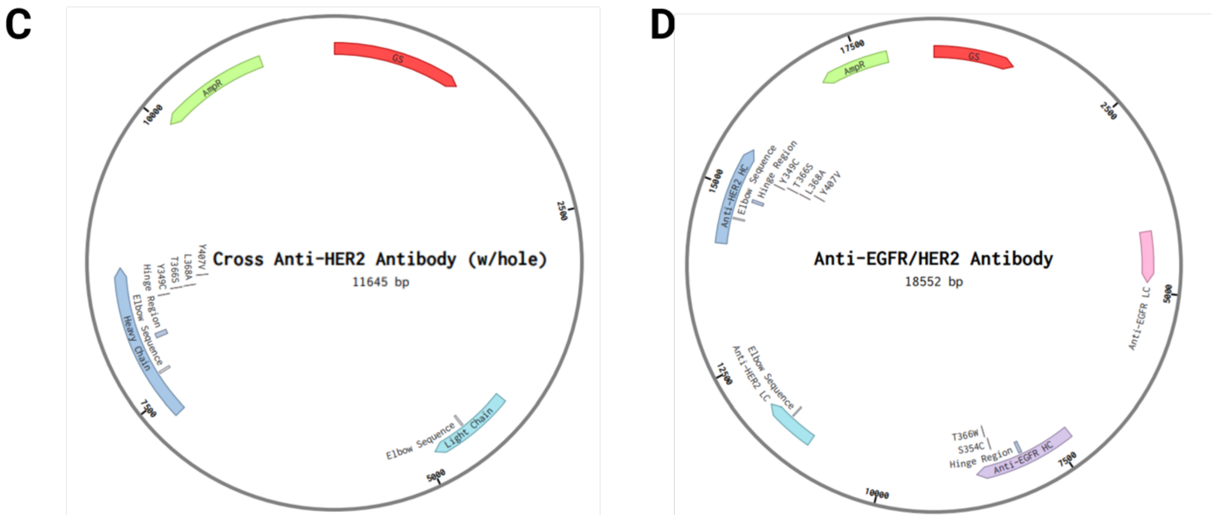


**Figure S2.** The representation of plasmid maps including monospecific and bispecific antibody expression cassette. (A-D) All recombinant proteins were expressed under the promoter CMV. Plasmid maps for the native monospecific antibody, knob-mutated and hole mutated versions of the monospecific antibodies used in this study, as well as the gene cassettes placed in a single plasmid for expressing bispecific antibodies, were designed on Benchling [1].

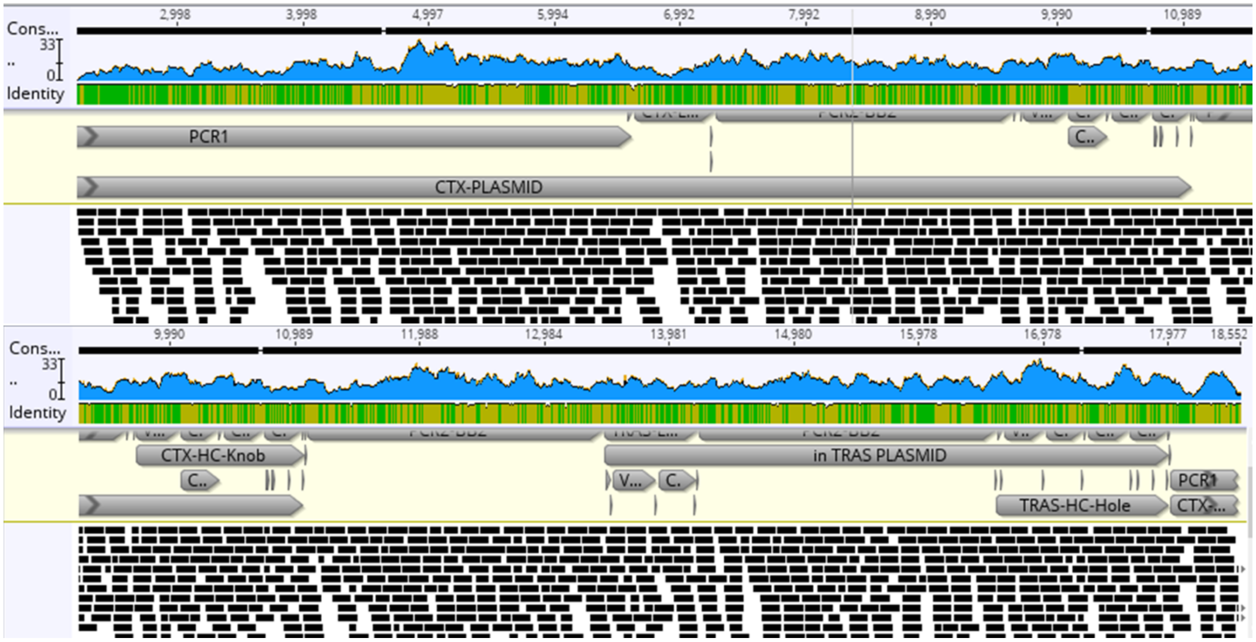


**Figure S3.** Next-generation sequencing analysis confirming cloning of the knob-into-hole–engineered Anti-EGFR/HER2 bispecific antibody.

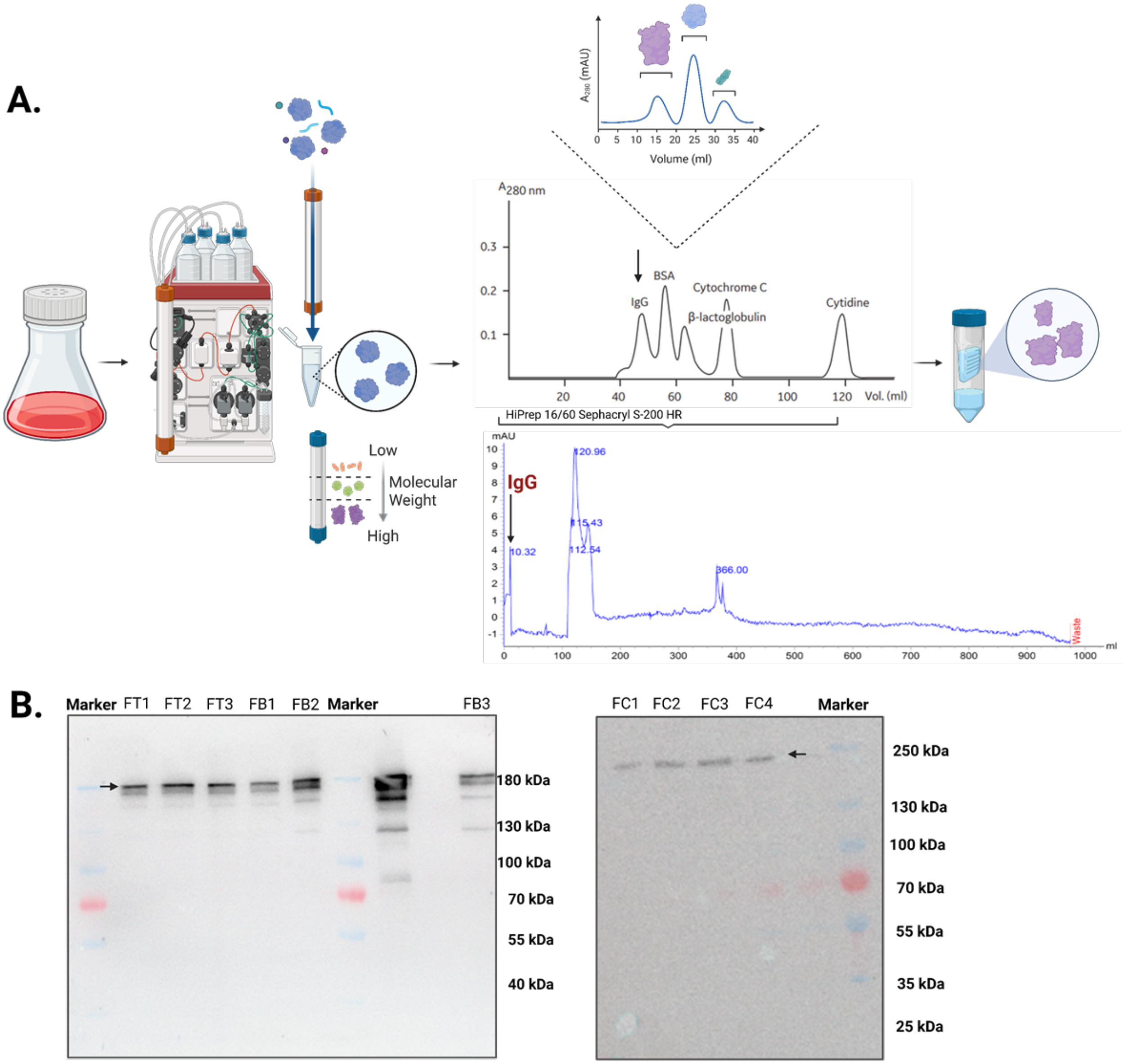


**Figure S4.** Samples were processed through a multi-step preparation and purification workflow utilizing fast protein liquid chromatography (FPLC). (A) Initially, cell suspensions were centrifuged, and the resulting supernatants were carefully collected. To prevent column clogging during purification, samples were filtered prior to loading. Antibody purification was subsequently carried out using the FPLC system. The resulting purified antibody fractions were then employed in downstream cell-based experiments. All protein samples were analyzed by Western blotting to confirm purity and integrity. The schematic representation was created using BioRender.com. (B) Western blot analysis was performed on the FPLC-purified fractions of Anti-EGFR, Anti-HER2, and bispecific Anti-EGFR/HER2 antibodies. All samples exhibited bands around ~190 kDa, consistent with the expected molecular weight of full-length IgG antibodies. Labeled as follows: FC – fractions collected in the Anti-EGFR elution tube; FT – fractions from the Anti-HER2 purification; FB – fractions corresponding to the bispecific Anti-EGFR/HER2 antibody.

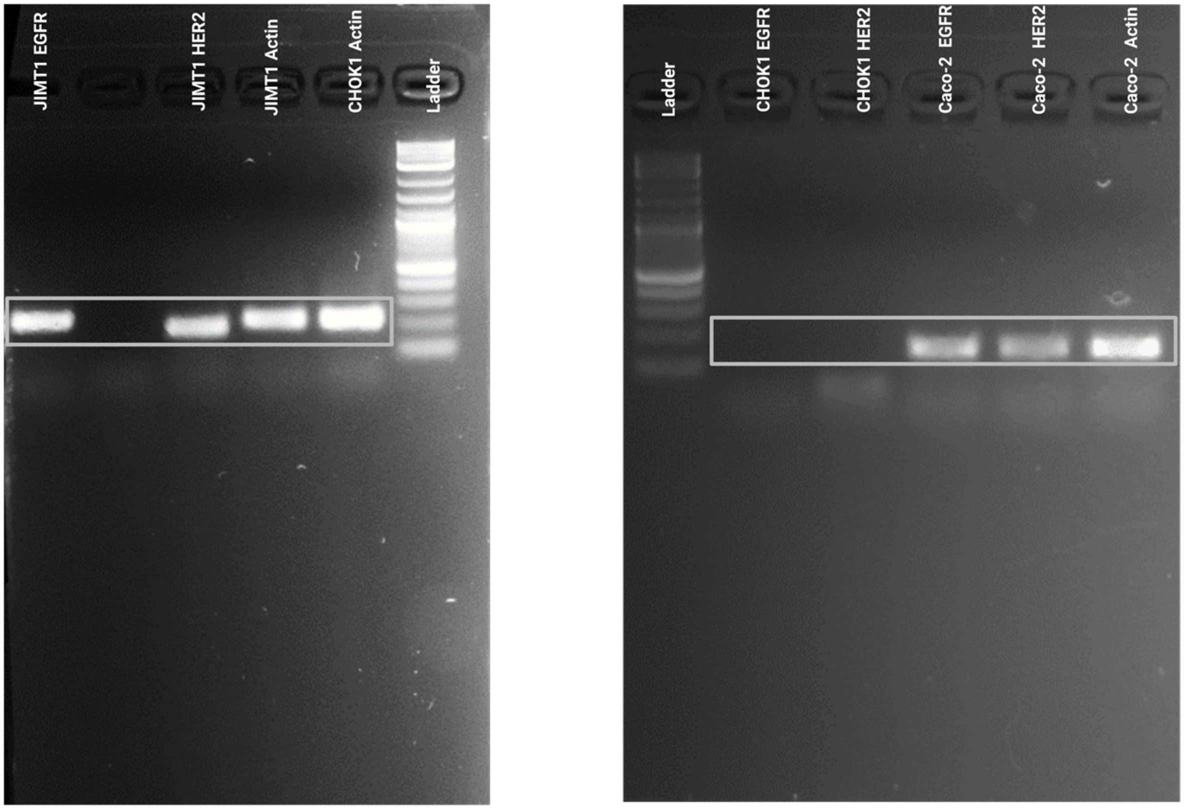


**Figure S5.** Bispecific antibodies were used to demonstrate their efficacy using JIMT-1 and Caco-2 cell lines. Following qPCR to confirm that both cell lines expressed EGFR and HER2, the products were run on an agarose gel. In the CHOK1 cells used for negative control, neither receptor was detected, while both antigens were expressed in the JIMT-1 and Caco-2 cell lines.

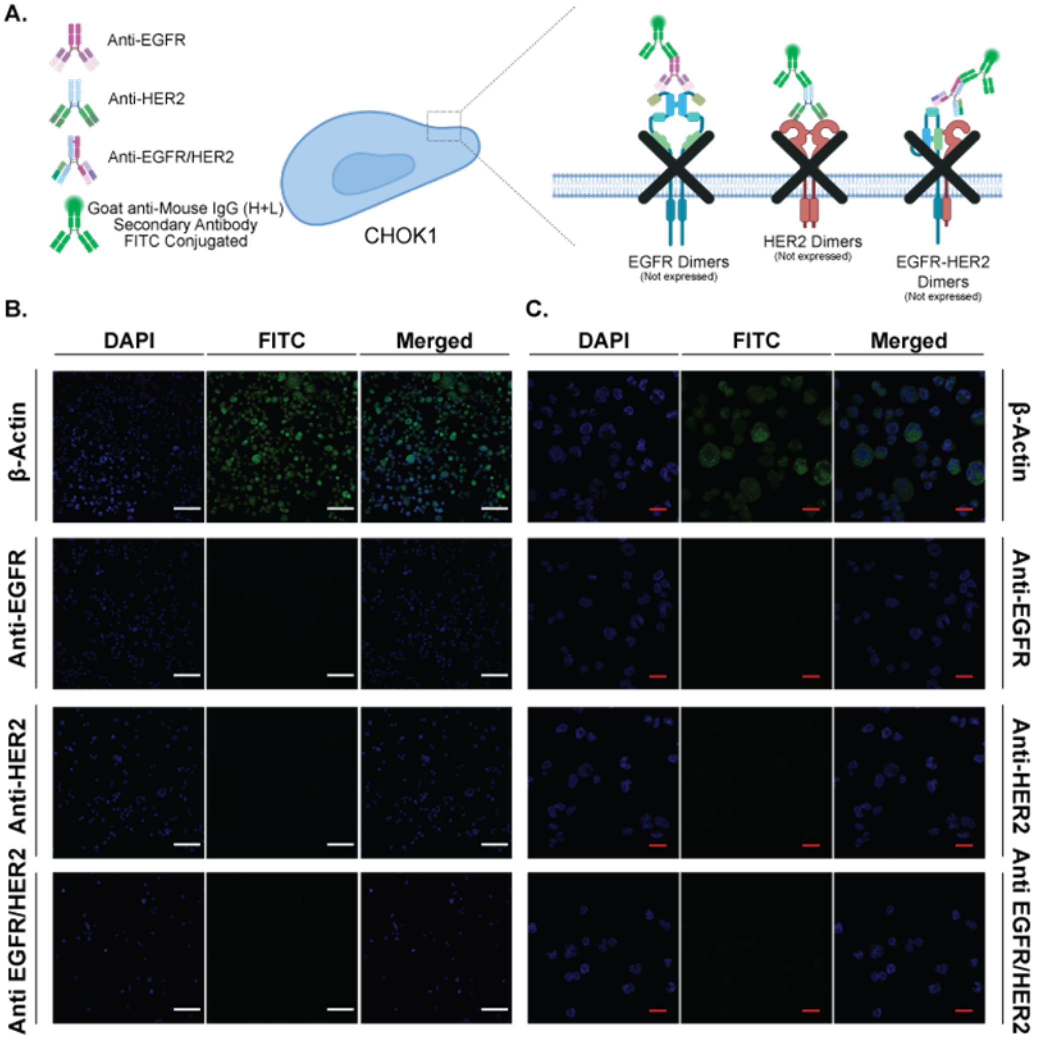


**Figure S6.** Immunocytochemistry analysis confirms the absence of EGFR and HER2 expression in CHOK1 cells, validating antibody specificity. A. Schematic drawing of anti-EGFR, anti-HER2, and bispecific anti-EGFR/HER2 antibodies and their conceptual binding on CHOK1 cells in the absence of endogenous EGFR and HER2 expression. (Created with BioRender.com) B. Confocal microscopy images of CHOK1 cells subjected to ICC. DAPI staining (blue) was used to visualize nuclei to confirm cell integrity. β-actin immunostaining was performed as a positive control which produced a clear FITC signal, demonstrating successful antibody detection and validating the ICC protocol. In contrast, ICC performed with anti-EGFR, anti-HER2, and bispecific anti-EGFR/HER2 antibodies generated no detectable FITC signal, consistent with the receptor-negative profile of CHOK1 cells. Such findings confirm the absence of EGFR/HER2 expression and demonstrate the specificity and absence of nonspecific binding of the engineered antibodies. White scale bars: 100 μm. Images were taken with 20X objectives. C. Confocal microscopy images of CHOK1 cells subjected to ICC. The images were obtained with 63X objectives. Red scale bars: 20 μm.

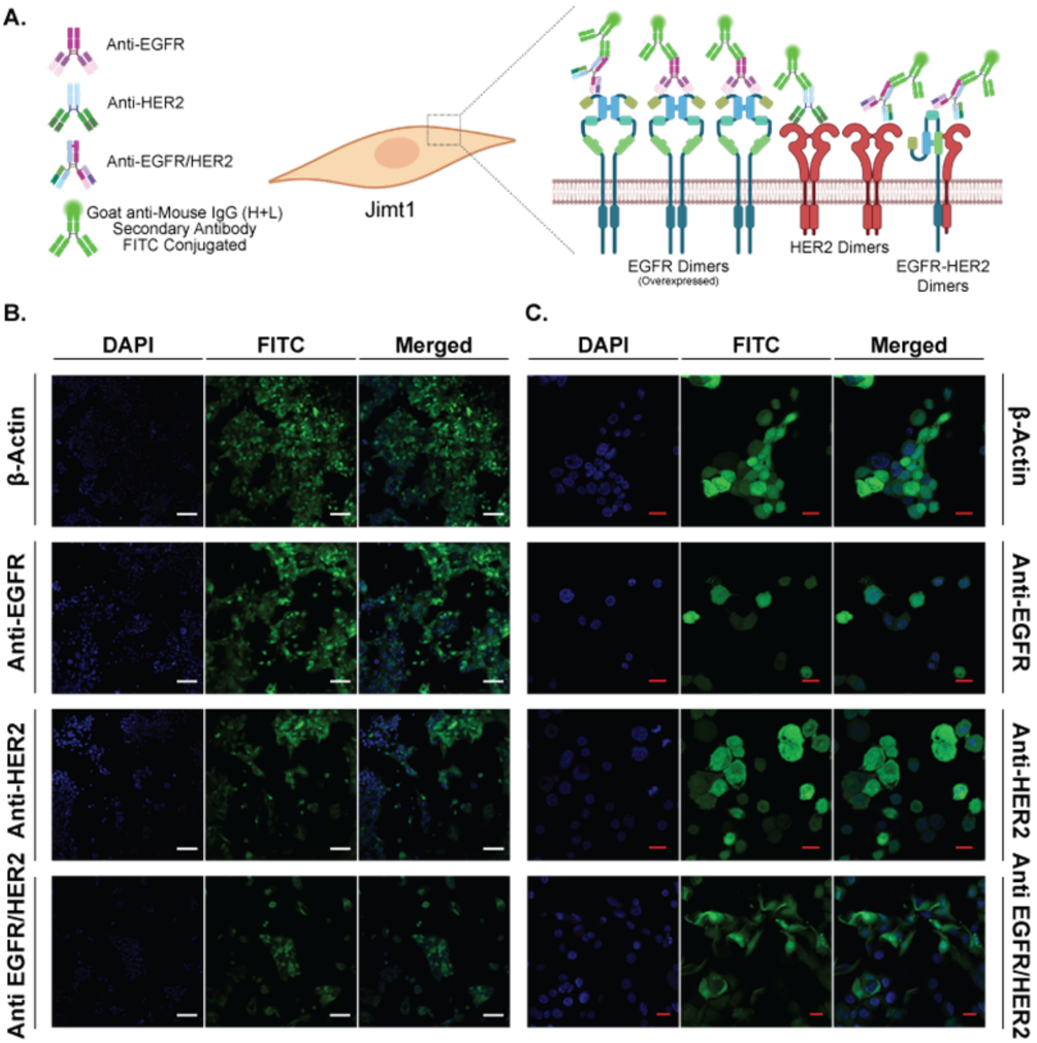


**Figure S7.** Immunocytochemistry analysis showing receptor-specific binding of anti-EGFR, anti-HER2, and bispecific anti-EGFR/HER2 antibodies in JIMT-1 cells. A. Schematic representation of the anticipated interactions of monospecific and bispecific antibodies with endogenous EGFR and HER2 receptors expressed on JIMT-1 cells. (Created with BioRender.com) B. Shown are confocal micrographs of JIMT-1 cells that were stained with DAPI (blue) and probed with anti-EGFR, anti-HER2, or bispecific anti-EGFR/HER2 antibodies, followed by FITC-conjugated secondary antibodies (green). As expected from this cell line that naturally expresses both EGFR and HER2, JIMT-1 cells showed distinct membrane-localized FITC fluorescence after all the various antibody treatments. In agreement with the receptor expression profile of this cell line, the highest FITC signal was obtained with anti-EGFR staining, reflecting the high abundance of EGFR, whereas anti-HER2 produced weaker yet distinct fluorescence. The bispecific antibody induced intense and diffuse fluorescence consistent with its dual-targeting capability. DAPI staining confirmed proper cellular integrity and nuclear localization. Scale bars: 100 µm C. Scale bar: 20 µm. The images of the same samples were obtained using a 63X objective.

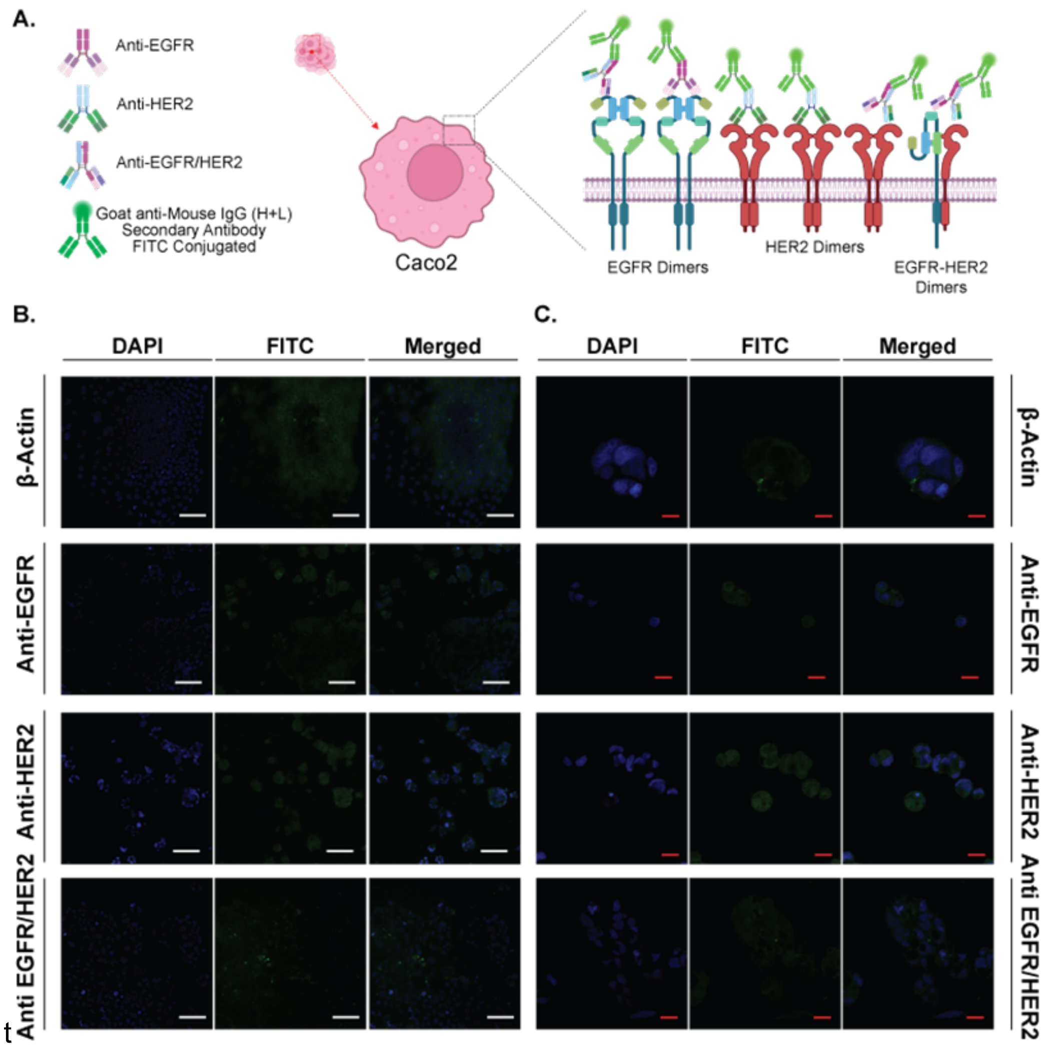


**Figure S8.** Immunocytochemistry analysis of the binding of anti-EGFR, anti-HER2, and bispecific anti-EGFR/HER2 antibodies in Caco-2 cells. A. Schematic overview illustrating antibody–receptor interactions on Caco-2 cells, which possess polarized epithelial architecture and tightly packed junctional complexes that hinder antibody access to surface receptors. (Created with BioRender.com) B. Representative confocal images of Caco-2 cells stained with DAPI (blue) and probed with anti-EGFR, anti-HER2, or anti-EGFR/HER2 bispecific antibodies, followed by FITC-conjugated secondary antibodies (green). Due to the aggregated, dome-forming morphology characteristic of Caco-2 cultures, FITC fluorescence was substantially weaker compared with JIMT-1 or CHOK1 cells. Limited staining is consistent with known challenges in antibody penetration across compact epithelial layers and restricted lateral receptor accessibility in polarized intestinal epithelial models. The mild FITC signals corresponding to anti-EGFR and bispecific antibody binding were observed in less dense regions, whereas anti-HER2 staining remained minimal, reflecting lower HER2 expression. DAPI staining confirms nuclear integrity across all samples. Scale bar: 100 µm C. The images were taken from the same samples with 63X objectives. Scale bar: 20 µm.

### Supplementary Material and Methods

For purification, HiTrap Protein A HP, 1 x 1 ml (29048576) and HiTrap MabSelect PrismA 1 x 1 mL (17549851) were used.

*Protein A Affinity Chromatography.* Roche cOmplete™ Protease Inhibitor Cocktail (11697498001) was added to the harvested samples, which were subsequently mixed 1:1 with binding buffer. Prior to sample loading, the column was equilibrated with 5–10 column volumes (CV) of binding buffer (20 mM sodium phosphate, pH 7.2). Samples were loaded at a flow rate of 0.5 mL/min. Elution was performed using 0.1 M citric acid buffer at pH 3.0 for monospecific antibodies and pH 4.5 for bispecific antibodies.

*MabSelect Affinity Chromatography.* Samples were supplemented with Roche cOmplete™ Protease Inhibitor Cocktail (11697498001) and mixed 1:1 with binding buffer prior to purification. The column was equilibrated with 5–10 CV of binding buffer (20 mM sodium phosphate, 0.15 M NaCl, pH 7.2), and samples were loaded at a flow rate of 0.5 mL/min. Elution was achieved using 0.1 M sodium citrate at pH 3.0 for monospecific antibodies and pH 3.5 for bispecific antibodies.

### Supplementary References

[1] Benchling [Biology Software]. Retrieved from https://benchling.com
